# Autophagy activators normalize aberrant Tau proteostasis and rescue synapses in human familial Alzheimer’s disease iPSC-derived cortical organoids

**DOI:** 10.1101/2025.06.25.661453

**Authors:** Sergio R. Labra, Jadon Compher, Akhil Prabhavalkar, Mireya Almaraz, Claudia Cedeño Kwong, Christine Baal, Maria Talantova, Nima Dolatabadi, Julian Piña-Sanz, Yubo Wang, Leonard Yoon, Swagata Ghatak, Zi Gao, Yuting Zhang, Dorit Trudler, Lynee Massey, Wei Lin, Anthony Balistreri, Michael Bula, Nicholas J. Schork, Tony S. Mondala, Steven R. Head, Jeffery W. Kelly, Stuart A. Lipton

## Abstract

Alzheimer’s disease (AD) is the most common form of dementia worldwide. Despite extensive progress, the cellular and molecular mechanisms of AD remain incompletely understood, partially due to inadequate disease models. To illuminate the earliest changes in hereditary (familial) Alzheimer’s disease, we developed an isogenic AD cerebrocortical organoid (CO) model. Our refined methodology produces COs containing excitatory and inhibitory neurons alongside glial cells, utilizing established isogenic wild-type and diseased human induced pluripotent stem cells (hiPSCs) carrying heterozygous familial AD mutations, namely PSEN1^ΔE9/WT^, PSEN1^M146V/WT^, or APP^swe/WT^. Our CO model reveals time-progressive accumulation of amyloid beta (Aβ) species, loss of monomeric Tau, and accumulation of aggregated high-molecular-weight (HMW) phospho(p)-Tau species. This is accompanied by neuronal hyperexcitability, as observed in early human AD cases on electroencephalography (EEG), and synapse loss. Single-cell RNA-sequencing analyses reveal significant differences in molecular abnormalities in excitatory vs. inhibitory neurons, helping explain AD clinical phenotypes. Finally, we show that chronic dosing with autophagy activators, including a novel CNS-penetrant mTOR inhibitor-independent drug candidate, normalizes pathologic accumulation of Aβ and HMW p-Tau, normalizes hyperexcitability, and rescues synaptic loss in COs. Collectively, our results demonstrate these COs are a useful human AD model suitable for assessing early features of familial AD etiology and for testing drug candidates that ameliorate or prevent molecular AD phenotypes.

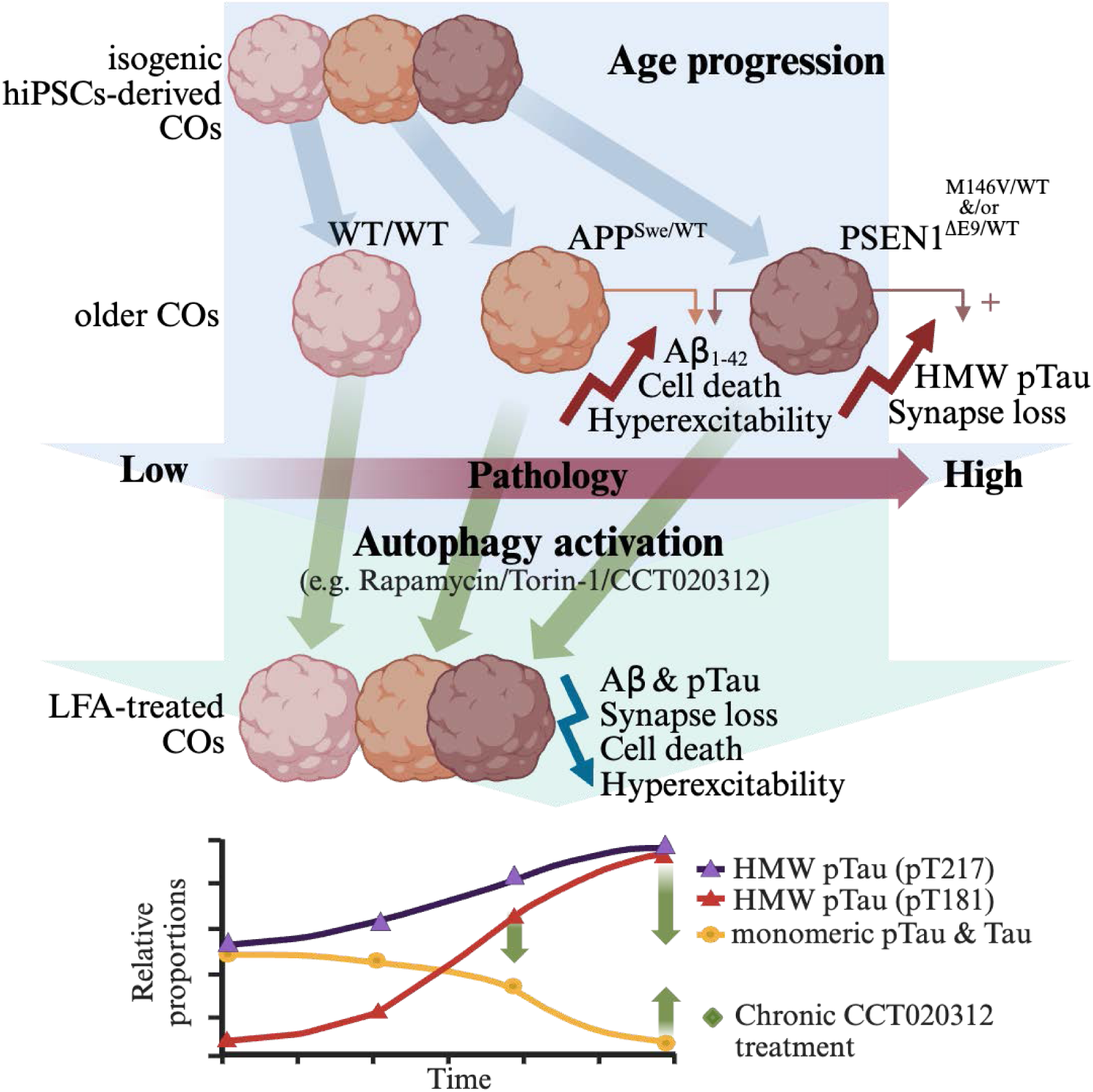

## INTRODUCTION

Alzheimer’s disease (AD) is a neurodegenerative disorder, currently affecting over 47 million people worldwide, characterized by synapse and neuronal loss leading to progressively declining cognitive function.^1,2^ The consensus view of AD pathology is that Ab aggregation, triggering hyperphosphorylated Tau aggregation, leads to neuronal dysfunction and ultimately brain cell death.^3^ These aggregates compromise membrane integrity and organelle function, while also altering transcriptional programs and engaging signaling pathways, leading to multiple mechanisms of pathobiology, including neuroinflammation.^2,4^ Most AD therapies targeting amyloid beta (Aβ) — including β-secretase or γ-secretase inhibitors and aggregation blockers — have failed in clinical trials due to limited efficacy or adverse effects. Recent exceptions are the FDA-approved anti-Aβ antibodies lecanemab and donanemab, which modestly slow progression of AD-associated mild cognitive impairment by 25-30%, in part via microglia-mediated lysosomal clearance of Aβ aggregates.^1^ In contrast, to date no clinically approved therapies effectively halt Tau pathology. Although early-stage active immunotherapy against pathological Tau protein shows limited promise in Aβ/Tau-positive patients, most have failed.^5,6^ Collectively, these clinical findings underscore Ab aggregation as a driver of AD progression, with Tau pathology and neuroinflammation as additional contributing factors.

The advent of human induced pluripotent stem cells (hiPSCs) derived from AD patients harboring mutations in amyloid precursor protein (APP) or presenilin 1 (PSEN1; one of four proteins in the g-secretase complex) and the generation of isogenic, gene-corrected controls afforded us the opportunity to capture the earliest changes associated with AD longitudinally in specific differentiated tissues of interest. While traditional two-dimensional (2D) cultures have numerous practical advantages, they fall short in encompassing the complexity of neural tissue^7,8^. Advances in 3D culture systems now enable the creation of self-organizing cerebrocortical organoids (COs). These COs recapitulate essential aspects of human brain development, including the development of multiple cellular identities like astrocytes and diverse neuronal classes, though notably lacking microglial cells^9–12^. Previous studies using hiPSC-derived neuronal and astrocytic 3D co-cultures from AD patients have shown the formation of Aβ aggregates and Tau neurofibrillary tangles (NFTs)—hallmark pathologies not easily observed in most animal or 2D culture models^13–17^. These pathological molecular phenotypes are attenuated or reversed by β-secretase and □-secretase inhibitor treatment, validating animal model findings and demonstrating the drug-testing potential of 3D culture models^17–19^. We reasoned that a CO-based AD model would allow for a more pathophysiologically faithful cellular function and response system characterization than 2D culture models, due to its intricate and diverse cellular organization, albeit for early events in disease pathogenesis. This strategy has been previously proposed by others, especially with regard to the potential utility for testing novel therapeutic candidates.^13–15,20–25^ Nonetheless, concerns still exist regarding the use of COs as models of intrinsically advanced age-related diseases like AD.^26–32^ It is currently unclear how these relatively young cellular models recapitulate phenotypes like NFTs that would normally take decades to develop in the human brain. Thus, we set out to develop and robustly validate an improved CO-based model to more faithfully represent the complex pathophysiological changes and cell-cell interactions in the early stages of familial AD (fAD). This model would facilitate the study of both, progressive responses to accumulating pathologic stress by different cell types and the potential causes behind this expedited pathology.

Previous studies have shown that hiPSC-derived neurons harboring the APP mutation recapitulate several key aspects of AD pathology, including elevated Aβ_1-42_or Aβ_1-42_/ Aβ_1-40_ ratio, increased Tau phosphorylation, and neuronal stress.^33,34^ However, these studies were limited by the use of 2D monocultures that lacked the complexity and cellular diversity of organoid cultures. Meanwhile, most studies on these PSEN1 mutations have been limited to clinical genetic or postmortem correlations with limited mechanistic insight,^35–40^ while the few in vitro studies performed to date have been limited in scope.^18,41–44^

Additionally, several lines of evidence suggest that the autophagy-lysosomal pathway is impaired in AD, contributing to the accumulation of toxic protein aggregates and neurodegeneration reflecting aberrant proteostasis.^45^ Postmortem brain tissues from AD patients exhibit an accumulation of autophagosomes and lysosomes, indicating a defect in autophagosome-lysosome fusion and lysosomal degradation.^46^ Moreover, familial AD-linked mutations in PSEN1 and amyloid APP have been shown to disrupt lysosomal acidification and proteolysis, leading to autophagic dysfunction. These findings have led to the hypothesis that enhancing autophagy and lysosomal function could be a promising therapeutic strategy for AD.^47,48^ Indeed, various autophagy activators, such as rapamycin, trehalose, and small molecule enhancers of rapamycin (SMERs), have been reported to reduce Aβ and Tau pathology to some degree in immortalized cell lines, as well as to alleviate cognitive deficits and slow disease progression in transgenic AD mouse models.^49–51^ However, the optimal dosing regimen and long-term efficacy of these compounds remain to be established. Here, we make initial strides to establish a CO dosing strategy employing mTOR inhibitor-independent autophagy activators. This is particularly important as the US Food and Drug Administration (FDA) has recently signaled that it may accept such data in lieu of and in preference to animal data for future drug approvals.^52,53^

In this study, we employed previously established gene-edited hiPSC lines carrying heterozygous fAD mutations in the genes encoding presenilin 1 (PSEN1) or amyloid precursor protein (APP) and compared results to their isogenic, wild-type (WT)-gene controls.^34,54^ Specifically, we used a PSEN1^M146V/WT^ missense mutation (M146V/WT), which enhances Aβ 1-42 peptide (Aβ_1-42_) production leading to an increased Aβ_1-42_/ Aβ_1-40_ ratio, a hallmark associated with an aggressive early-onset AD phenotype. Additionally, the APP^Swe/WT^ double mutation (K670N/M671L) residing directly upstream of the β-secretase cleavage site, makes APP a more efficient substrate, leading to increased total Aβ production. Together, these mutants comprise our isogenic set 1.^55–57^ To contrast with a different genetic background,^58^ we also utilized a line with the PSEN1^ΔE9/WT^ mutation (delE9/WT), an in-frame deletion of exon 9 (residues 291-319) that leads to impaired endoproteolytic processing of PSEN1, which also known to result in an increased Aβ_1-42_/ Aβ_1-40_ ratio.^35,59^ This mutation was introduced into a separate, well-established hiPSC line (with delE9/WT and WT/WT (2) cell lines comprising isogenic set 2).^41,60^

Herein, we report the development of a thoroughly characterized human AD CO model based on hereditary AD mutants and isogenic controls. We describe a protocol that reproducibly generates COs and show that the COs exhibit the hallmark biochemical, transcriptomic, and functional abnormalities of AD. Furthermore, we demonstrate their use to interrogate, in real time, the effects of using autophagic induction as a therapeutic strategy.

## RESULTS

### Development of a consistent and robust AD CO model

To consistently produce comparable COs across all our cell line genotypes, we started with the elegant method published by Paşca and colleagues, which we designate Protocol 1.^3^ We optimized Protocol 1 to minimize CO fusion, prevent CO attachment to the surface, reduce self-disassembly, and improve viability in our COs, generating protocol 2 (**Figures 1A** and **S1A**). Maintaining the COs under constant mixing in ULA 6-well plates by employing an in-incubator rocker and/or tilted orbital shaker (15° incline, 65 rpm) significantly improved CO viability and morphology (**Figure S1B**). Staining live COs with propidium iodide at around the 4-month timepoint showed that necrosis was also reduced, suggesting improved nutrient absorption within the COs (**Figure S1C**). Next, we studied the morphology and size distribution of COs produced by modifying the induction medium. We compared Protocol 1’s Gibco’s Essential 6 medium (E6) versus different concentrations of hESC medium, given the latter’s use in other CO induction methods.^10,61^ We found that a mixture of 50% E6 and 50% hESC medium (“E6/hESC”) yielded larger COs with more pronounced neuroectoderm morphology than those grown in E6 alone (**Figures S1D-F**), designated herein as Protocol 2. Importantly, Protocol 2 with the E6/hESC medium produced COs without significant size differences across all the different genotypes by the 5-week time point in culture (**Figure 1B** and **S1G**). Despite the COs consistently growing larger in size, they showed dramatically lower levels of necrosis compared to those grown in parallel following protocol 1 (**Figure S1C**).

**Figure 1:**
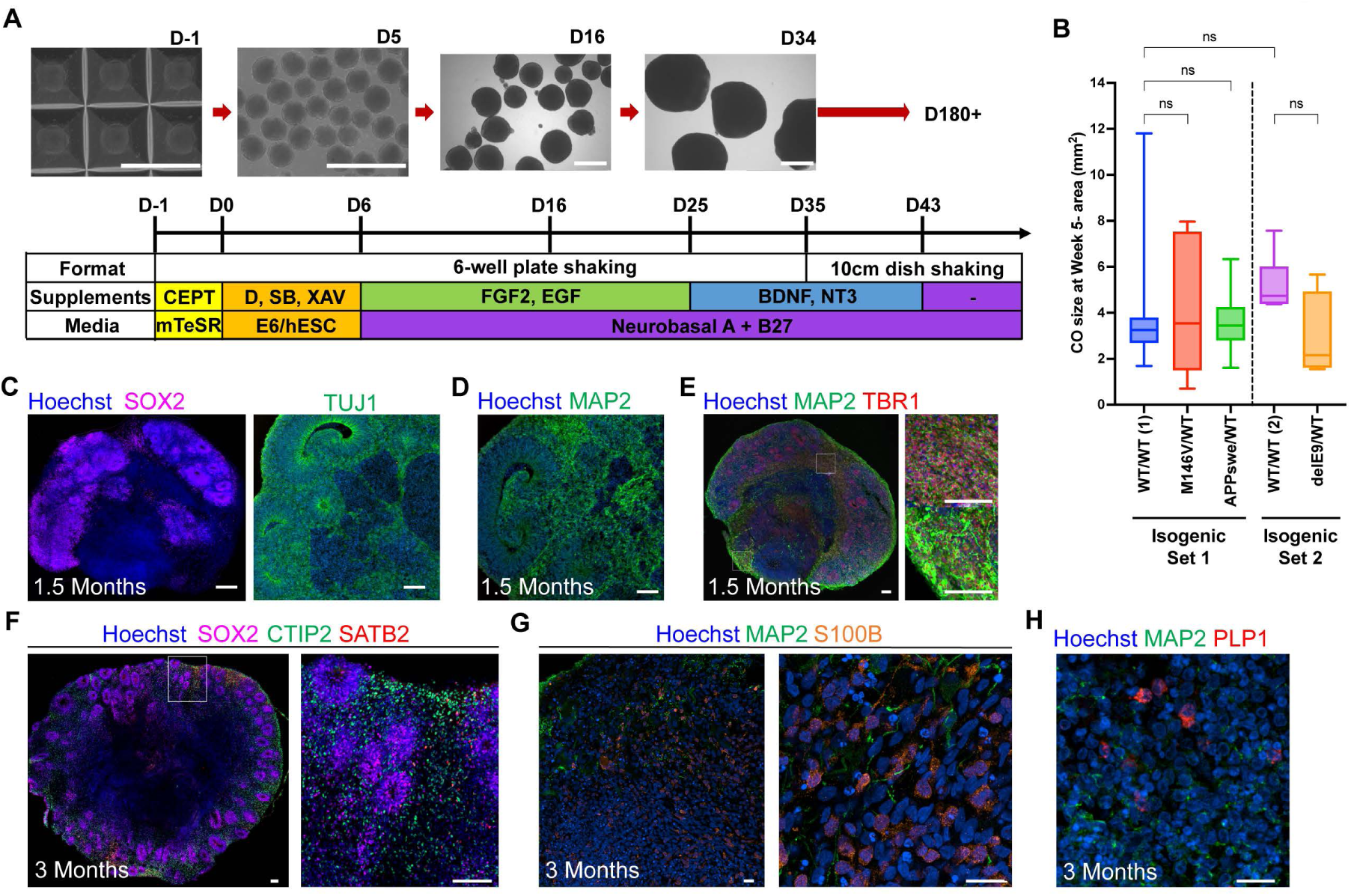
Generation of isogenic AD COs. (A) Schematic describing the differentiation of cerebrocortical organoids (COs) with representative brightfield images. Scale bar, 1 mm. (B) Quantification of the CO maximal cross-sectional area distribution across genotypes at approximately week 5 in culture, n = 13-26 COs per genotype for isogenic Set 1, n = 6-9 COs per genotype; each from 4 independent CO induction batch experiments. Data are mean ± SD. Analysis by ANOVA with Dunnett’s post-hoc test. (C) Representative immunofluorescence (IF) stains of neuroepithelial (SOX2^+^) and neuronal progenitor (TUJ1^+^) cells at ∼7-week timepoint COs. Scale bar, 100 µm. (D) Representative IF stains of mature postmitotic neurons (MAP2^+^) in COs at ∼7-week timepoint. Scale bar, 100 µm. (E) Representative IF stains for mature cortical layer neurons (TBR1^+^) cells in a 7-week timepoint CO with two highlighted insets to the right. Scale bar, 100 µm. (F) Representative IF stains for deep layer V (CTIP2^+^) and upper layer II-III (SATB2^+^) cells in a 3-month timepoint CO with inset to the right. Scale bars, 100 µm. (G) Representative IF stains of astrocytes (S100β^+^) 3-month timepoint COs. Scale bars, 25 µm. (H) Representative IF stains of oligodendrocytes (PLP1^+^) 3-month timepoint COs. Scale bar, 25 µm. See also Figure S1.

To scrutinize the differentiation quality of the COs, we performed immunohistochemical labeling of fixed, sliced CO samples. At day (D)50, SOX2 and TUJ1 staining demonstrated the presence of neural progenitors most densely accumulated around ventricular-like rosette formations (**Figure 1C**). Meanwhile, MAP2^+^ mature neurons populated the large majority of the COs outside the margin of these rosettes (**Figure 1D**); NeuN was also used as a mature neuron marker (**Figures S1H**). Staining for FOXG1, a critical forebrain transcription factor key in cerebrocortical development, showed the organoids to be maturing toward forebrain identity (**Figure S1I**). Furthermore, TBR1 staining revealed a large number of MAP2^+^/TBR1^+^ number of cells, maximizing their abundance throughout the variable structure folding throughout the CO (**Figure 1E**). Staining neurons in various cerebrocortical layers using layer-specific markers, including SATB2 (upper cortical layers 2-4) and CTIP2 (deep cortical layers 5-6), showed non-overlapping positive signals spatially distributed across the COs around the SOX^+^ rosettes (**Figure 1F**). Together, these findings strongly suggest that the cells within the developing COs emulate ventricular zone migration, and that the various neuronal layer subtypes are properly differentiating without acquiring overlapping subtype identities. Previously, this has been highlighted as a concern in other CO differentiation protocols.^26^

Staining for astrocytes via S100B (**Figure 1G**) demonstrated populations of astrocytes are already abundant in the COs at the 3-month timepoint. Notably, we also found the presence of PLP1+ cells, putatively oligodendrocytes, (**Figure 1H**) by the 3-month timepoint. The presence of these cells is particularly striking given that previous CO-based studies have struggled to generate any or enough of these critical glial cells.^62^

To further characterize the cellular diversity and reproducibility of our COs, we performed single-cell RNA sequencing (scRNA-seq) at two timepoints, after ∼7-8 weeks and 3 months in culture. We multiplexed isogenic WT and fAD COs differentiated in parallel using the 10x Genomics platform (Illumina) at ∼40K reads per sample across multiple independent experiments. Each sequencing dataset underwent separate, stringent quality control before integrating and reprocessing for annotation and validation (**Figure S2A**; **STAR Methods**). The final dataset consisted of 70,854 cells. Overall, there was significant overlap between the two timepoints (**Figure S2B**), and each AD and WT genotype contributed to all unsupervised clusters (**Figures S2C** and **S2D**), indicating the clustering was not significantly driven by genotype-specific differences. We identified ten major cell-type groups (**Figure 2A**). The localized expression of key hallmark markers showed the specificity of each validated cell-type (**Figures 2B** and **S2E**).^63–67^

**Figure 2:**
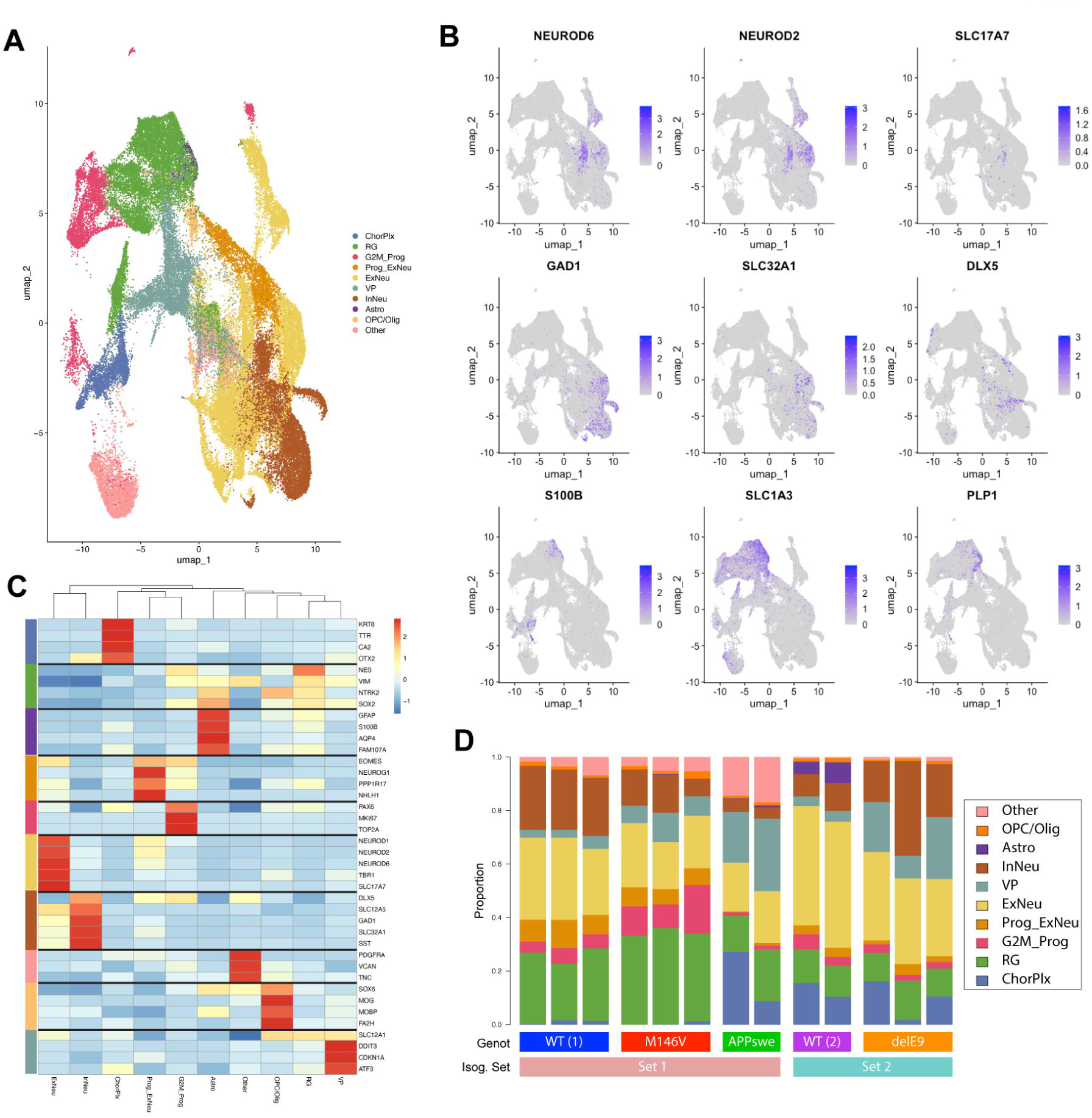
COs recapitulate key cellular diversity. (A) Annotated UMAP clustering of CO’s single cell-RNA-seq at 2-month and 3-month timepoints, n = 27 COs in total from 5 independent experiments, processed into 4 sequencing runs. (B) Feature plots demonstrating the specific segregation of cell-types by the hallmark gene marker expression of key cell type markers. Highlighted here are Excitatory neurons (NEUROD6 and SLC17A6, aka VGLUT2); Inhibitory neurons (GAD1 and SLC32A1); Astroglia (S100B), and Oligodendrocytes (PLP1). (C) Heatmap ordered by similarity showing the differential expression of hallmark marker genes in all identified cell subtypes. See also Figure S2F. (D) Bar plot showing the relative composition or abundance of cell types in the COs at 3-month timepoint. See also Figures S2, S3, S4, and S5.

As examples, expression of NEUROD6, NEUROD2, and SLC17A7 reflected excitatory neurons; GAD1, SLC32A1, and DLX5, inhibitory interneurons; SLC1A3, glial cells; S100B, astrocytes; and PLP1, oligodendrocytes, specifically. Cell-type similarity analysis agreed with the expression and relatively low overlap in key hallmark genes across the different cell types, suggesting proper differentiation and maturation (**Figure 2C** and **S2F**). Of particular interest, the presence of oligodendrocytes was also validated by their unique expression of additional markers like MOG, MOBP, and FA2H. We also note that across all the samples, the COs are most similar to COs sharing their own genotype with minimal variation between them, showcasing consistency (**Figures 2D** and **S2G**). These results suggest our COs reproducibly developed mature cortex-relevant cell types, including excitatory and inhibitory neurons and their progenitors, astroglia, oligodendrocyte progenitor cells (OPCs), and oligodendrocytes.

### AD mutation drives stress-and senescence-related cell phenotypes in CO and abnormal neuronal development

We note that the VM-, SOX2-, and SLC12A1-positive CO ventral progenitors (VP) found through scRNA-seq were highly enriched in markers of cellular stress, including DDIT3, CDKN1A (p21), and ATF3 (**Figure 2C**).^68–72^ These “stressed” progenitors were significantly more abundant in the PSEN1^M146V/WT^, APP^Swe/WT^ and PSEN1^DE9/WT^ COs compared to their isogenic WT controls (**Figure S2H)**. Other differences included a significantly reduced proportion of interneurons (InNeu) in the PSEN1^M146V/WT^ and APP^Swe/WT^ COs and reduced excitatory neurons (ExNeu) in all AD mutant COs compared to their isogenic WT COs **(Figure S2H**).

We confirmed these scRNA-seq results through immunolabeling on fixed 6 week and 3 month CO slices for vesicular glutamate transporter 1 (VGLUT1) and g-aminobutyric acid (GABA) to estimate the populations of excitatory and inhibitory neurons, respectively (**Figure S3A and S3B**). Their relative abundance generally agreed with the scRNA-seq results; namely, we confirmed an age-progressing relative reduction of ExNeu in the APP^Swe/WT^ and PSEN1^DE9/WT^ COs (**Figure S3C**), and a decrease in InNeu in the PSEN1^M146V/WT^ and APP^Swe/WT^ COs vs. WT (**Figure S3D**). The relative abundance of these different neuronal cells is critical given the importance of excitatory-to-inhibitory imbalance in AD, a pathologic feature we reported previously in a different CO model.^43,73,74^.

Next, we performed pseudotime analysis on the whole dataset using the Monocle3 toolkit.^75^ It re-clustered the cells into several macro-clusters, where the major cluster with highest pseudotime values was composed of ramifying sub-clusters of ExNeu or InNeu populations (**Figures S4A** and **S4B**). Notably, multiple branched neuronal populations stemmed from mostly single AD mutant genotypes. In particular, APP^Swe/WT^ was highly enriched in specific ExNeu branches while the PSEN1^DE9/WT^ was highly enriched at the end of both the ExNeu and InNeu trajectories, with the PSEN1^M146V/WT^ mutant just behind it (**Figure S4C**). This trajectory analysis suggested the AD COs’ neurons were developing/reaching additional neuronal states that their isogenic WT organoids were not, and these states were specific to the type of mutation involved (PSEN1 or APP). Furthermore, when looking at the pseudo-time expression of hallmark ExNeu and InNeu markers, we find the expected progressive increase and slight decline in the immature ExNeu marker NEUROD2, as it gives way to maturing ExNeu, as marked by SLC17A7 (**Figure S4D, *left***). However, looking at InNeu progression, we noticed a sharp decline in both immature (DLX2) and mature (GAD2) InNeu marker expression (**Figure S4E, *left***). Repeating the pseudo-time analysis without the AD mutants (WT genotypes only) revealed not only a very clear trajectory from radial glia progenitors diverging into glial and neuronal cell types (**Figures S4F-H**), but the WT pseudo-time neuron marker expression lacked the sharp decline in DLX2 and GAD2 expression (**Figure S4E, *right***). There was no notable difference between the two ExNeu marker expression pseudo-time analyses (**Figure S4D, *right***). This demonstrates the decrease in InNeu marker expression is driven by the AD mutation. Together, this suggests that AD mutants may be developing into abnormal or vulnerable neurons as they age—which has been recently suggested for the APPswe mutation^76^—and in particular, they are affecting inhibitory neurons.

Next, we performed differential gene ontology (GO) enrichment analyses of the excitatory and inhibitory neurons in each of the AD mutant COs, compared to their isogenic WT CO controls (**Figures S5A-F**). Interestingly, we found both PSEN1 mutations leading to deficits in numerous synapse-related pathways in excitatory neurons (**Figures S5A** and **S5B**), while synapse-related terms were upregulated in the inhibitory neurons (**Figures S5D** and **S5E**). Similarly, the excitatory neurons of the PSEN1^M146V/WT^ and PSEN1^DE9/WT^ had significantly upregulated genes related to cytoplasmic RNA translation and ribosome biogenesis; and axon guidance and development, respectively (**Figures S5A** and **S5B**). The same pathways were depressed in both, PSEN1^DE9/WT^ and APP^Swe/WT^ inhibitory neuron mutants (**Figures S5E** and **S5F**). Finally, the APP^Swe/WT^ excitatory neurons manifest different upregulated pathways; In particular, protein folding-related stress responses were upregulated in excitatory neurons while RNA processing was upregulated in inhibitory neurons (**Figures S5C** and **S5F)**. These findings suggest these AD mutations affect starkly differently excitatory neurons from inhibitory neurons, which may in part explain the underlying development of excitatory/inhibitory (E/I) imbalance in AD.

The AD mutant COs were also found to generally have greater expression of senescence markers including p21 (CDKN1A), p16 (CDKN2A), and the senescence-associated secretory phenotype (SASP);^71,72,77^ this overexpression was weakly higher for the PSEN1^DE9/WT^, greater for the PSEN1^M146V/WT^, and strikingly increased in the APP^Swe/WT^ mutant compared to their respective isogenic WT controls (**Figure S5H**). Additionally, comparing across genotypes, we found that markers of ferroptosis^78,79^ are notably elevated in the PSEN1^M146V/WT^ (but not PSEN1^DE9/WT^) and APP^Swe/WT^ COs vs. WT (**Figure S5I**).

### AD COs recapitulate amyloid-b and phospho-Tau signatures of human AD brain

To characterize the extent to which the AD CO model recapitulates hallmark AD pathology, we measured the level of Aβ_1-40_ and Aβ_1-42_ peptides secreted into the conditioned media by the COs (**Figure 3A**). Using a clinical-grade Aβ peptide multiplex kit, we found that the APP^Swe/WT^ COs produce and secrete significantly more Aβ_1-40_ and Aβ_1-42_, while maintaining a similar Aβ_1-42_/Aβ_1-40_ ratio as the WT/WT (1) COs—perhaps not surprising since this is a mutation in APP rather than in the processing enzymes (**Figure 3B**). On the other hand, the PSEN1^M146V/WT^ mutant secreted relatively less Aβ_1-40_ compared to Aβ_1-42_, thus shifting the peptide products towards a significantly higher Aβ_1-42_/Aβ_1-40_ ratio (**Figures 3A** and **3B**). Notably, following the relative secreted Aβ_1-42_/Aβ_1-40_ ratio in COs from 3 to 7 months of age, we found the APP^Swe/WT^ maintained a stable Aβ_1-42_/Aβ_1-40_ ratio while the elevated PSEN1^M146V/WT^ mutant’s ratio progressively decreased over time (**Figure S6A**). This is remarkably the same progression numerous mass-spectrometry-validated studies have found in CSF of patients with fAD mutations compared to non-carrier controls decades before symptom onset.^80–82^ Isogenic set 2 revealed that PSEN1^DE9/WT^ COs had increased Aβ_1-42_levels as well as an increased Aβ_1-42_/Aβ_1-40_ ratio, similar to PSEN1^M146V/WT^ mutant COs. This ratio was also not significantly different between the two different WT genotypes. Finally, Aβ_1-38_ peptide measurements demonstrated more than a two-fold reduction in the PSEN^M146V/WT^ mutant compared to WT, but a moderate increase in the APP^Swe/WT^ mutant (**Figure S6B**). This finding agrees with previous work on PSEN1 and could suggest an inverse relationship between the production of Aβ_1-38_ and Aβ_1-42_.^83–85^ Aβ_1-38_ was below the detection limit in Isogenic Set 2. Increased abundance was confirmed via IF in all fAD mutant COs, even identifying strong amyloid signal in 9 months of age COs with the use of Amytracker (**Figure S6C**).^86,87^

**Figure 3:**
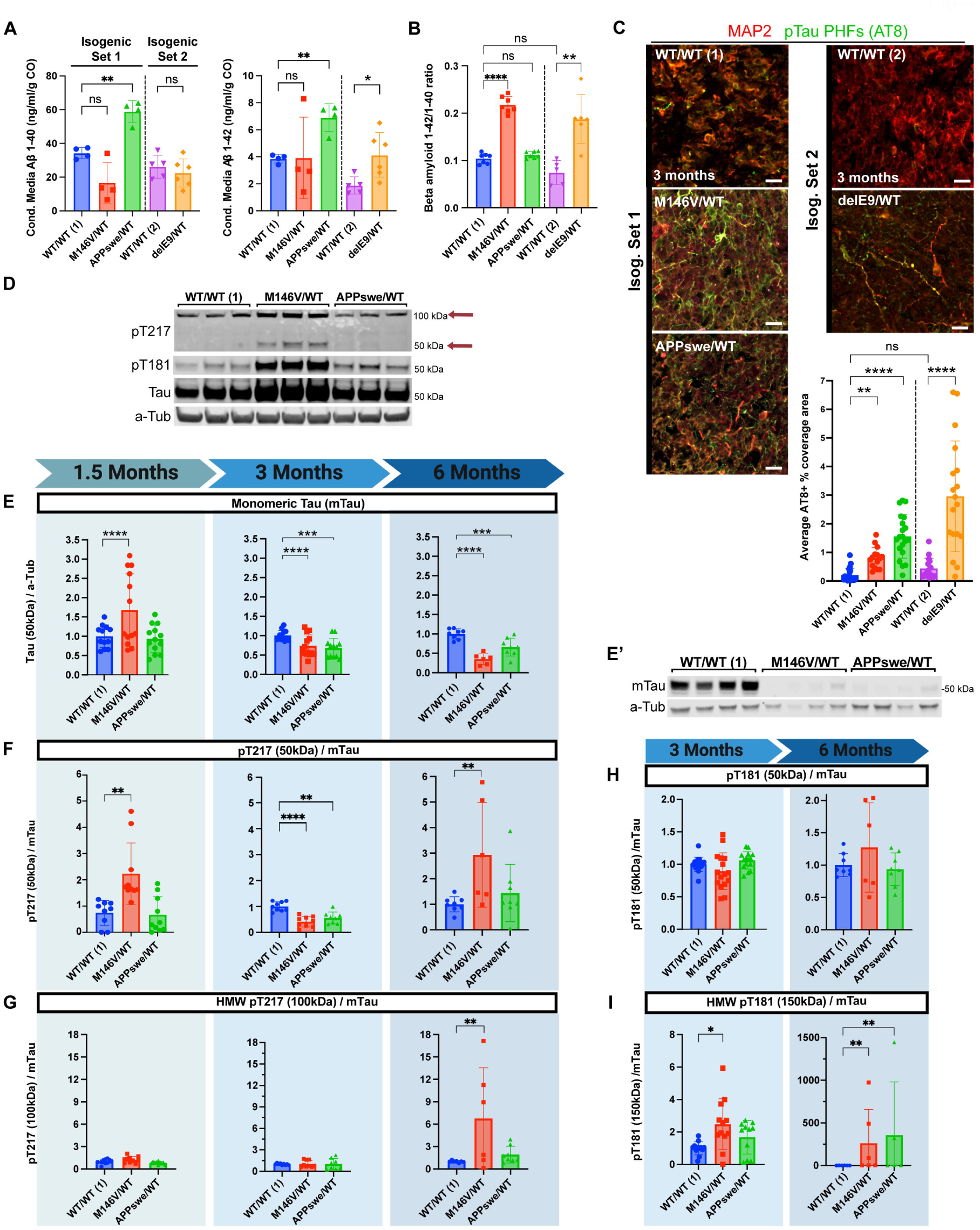
AD COs recapitulate amyloid and pTau pathologic signatures. (A) Aβ_1-40_ and Aβ_1-42_ concentrations in conditioned media from 5-6 months-of-age COs. n = 4-5 independent CO replicates per genotype. (B) Ratio of Aβ_1-42_ to Aβ_1-40_ from panel A. (C) Immunostaining of 3 months-of-age COs for Paired Helical Filament (PHF)-Tau (AT8, green) and MAP2 (mature neurons, red). Scale bar, 20 µm; alongside the final quantification. Isog. Set 1: n = 10-18 COs per genotype from 3 independent experiments; Isog. Set 2: n = 10-11 COs per genotype from 2 independent experiments. (D) Representative western blots (WB) of CO lysates at 1.5-month timepoint for phosphorylated Tau pT217, pT181 at its MS-validated monomeric molecular weight, pan-Tau at monomeric molecular weight (mTau hereafter), and -Tubulin (a-Tub). (E) WB Quantification of monomeric Tau (mTau) in CO lysates at the 1.5-(*left*), 3-(*center*), and 6-month (*right*) timepoints, as normalized to -Tubulin (a-Tub). n = 8-14 COs per genotype from 3 independent experiments. (E’) Representative WB of mTau and a-Tub in late-stage (4.5-month timepoint) CO lysates. (F-G) WB Quantification of phospho-Tau pT217 at its monomeric (∼50 kDa)(F), and strongest high-molecular-weight (HMW)(∼100 kDa)(G) bands in CO lysates at the 1.5, 3, and 6-month timepoints, as normalized to mTau. n = 5-12 COs per genotype from 3 independent experiments per timepoint. (H-I) WB Quantification of the pTau T181 bands at monomeric (∼50 kDa) (K) and strongest HMW (∼150 kDa) (L) bands at 3- and 6-month timepoints normalized to mTau. Data are mean ± SD. Analyses by ANOVA with Dunnett’s post-hoc test. See also Figures S6 and S7.

Given the strong link between Aβ and subsequent phospho-Tau pathologies, we employed the AT8 monoclonal antibody (recognizing Tau phosphorylated at Ser202 and Thr205), and commonly used to label pathologic paired helical filaments (PHF) via IF to probe the COs for pathological phosphorylated Tau (pTau).^88^ While both WT/WT genotypes had insignificantly different PHF-positive signal, we observed significantly increased PHF-positive staining in all the AD-associated mutants compared to their isogenic WT controls as early as 3 months of age (**Figure 3C**). Given this robust phenotype, we further investigated whether the COs recapitulated a progressive accumulation of clinically-relevant pTau as found in the human AD brain.^89^

We decided to characterize in the COs the two most clinically relevant pTau modifications, phospho-Thr217 (pT217) and phospho-Thr181 (pT181), which most closely correlate with disease progression in in CSF and plasma (particularly pT217).^90–93^ Although we attempted these measurements in conditioned medium, their abundance levels were found to be too low for detection via western blotting. However, we found both of these pTau species to be orders of magnitude more abundant in CO lysates. To control for the known limitation of “pan-Tau” or “total Tau” antibodies lacking sensitivity to many phosphorylated forms of Tau due to phosphorylation partially inhibiting their epitope binding,^94,95^ we initially focused our Tau quantification on the major band at the predicted monomeric molecular weight, denoted mTau hereafter (**Figures S6D** and **S6G)**.

Notably, we consistently found the appearance of specific high-molecular-weight (HMW) pTau^+^ bands, generally most abundant in the PSEN1^M146V/WT^ mutant COs (**Figure 3D**). We set out to rule out non-specific antibody binding. Using different antibodies, we found that apart from the expected monomeric bands around 50-55 kDa, the most abundant pT217^+^ band appeared at a MW consistent with dimers, at around 105 kDa (**Figures S6D-F)**.; while pT181 bands would often show a band at around 155 kDa, suggestive of trimers (albeit often too weak to quantify) (**Figures S6G-I)**. A limited number of previous clinical and animal studies have reported “low-n oligomeric” and “multimeric” pTau species detected via WB at approximately these MWs.^96–98^ Given the stringent reducing and solvating conditions of SDS-PAGE, the detection of these HMW species would suggest that they may be SDS-resistant and/or cross-linked in nature, suggesting these phospho-sites may be associated with oligomerization. To definitely validate these multimeric HMW bands, we performed immunoprecipitation (IP) enrichment of fresh PSEN1^M146V/WT^ COs and analyzed the protein content at these HMW bands via mass spectrometry (MS). Despite these HMW species not immunostaining with pan-Tau antibody (**Figure S6D** and **S6G**), MS confirmed Tau and pT217 IP-enriched bands at ∼50, ∼100, and ∼155 kDa, as they contained microtubule-associated protein Tau (MAPT) (**Figure S6J**, **STAR Methods**). Both pT181 and pT217 were identified, and WB highlighted that immunoprecipitated Tau contained significantly more pT181 than pT217. Thus, we quantified the respective HMW species independently from the monomeric ones.

Accordingly, we characterized Tau in CO lysates at the 1.5, 3, and 6-month timepoints. Consistent with the immunofluorescence (IF) data, we found mTau to be significantly elevated in the PSEN1^M146V/WT^ mutant COs at the 1.5-month timepoint (**Figures 3D** and **3E**). PSEN1 mutations have been associated with accelerated neural development,^18^ which may help explain this finding. However, we found that mTau progressively decreased in these PSEN1^M146V/WT^ COs over time, falling significantly lower at the 3-month timepoint, and further decreasing at later stages and below half of WT levels by 6 months of age (**Figures 3E** and **3E’**)— possibly reflecting the conversion of mTau into oligomers by this time point, thereby reflecting the disease process. Similarly, the APP^SWE/WT^ COs also demonstrated significantly decreasing mTau compared to WT COs after 3 months in culture, albeit with a less dramatic decline than the PSEN1^M146V/WT^ COs. This loss of mTau is also consistent with the notion that the AD COs exhibit microtubule destabilization, characteristic of the neurodegenerative process.^99^

Strikingly, the relative abundance of monomeric and HMW pT217 species manifested nearly opposite trends to one another in the AD COs compared to their isogenic WT controls (**Figures 3F** and **3G**). Early on, at 1.5 months of age, the PSEN1^M146V/WT^ COs displayed significantly increased monomeric pT217, while the no detectable differences in HMW pT217. As the organoids aged, this pattern shifted: monomeric pT217 levels decreased in this mutant by 3 months, while the HMW pT217 levels trended higher, reaching pronounced fold-changes by 6 months. Notably, this shift was even clearer in the second isogenic set’s PSEN1^DE9/WT^ COs. While they maintained relatively stable monomeric Tau levels (**Figure S6L**), these COs changed from only exhibiting significantly higher monomeric pT217 levels at 1.5-month timepoint, to developing significantly elevated HMW pT217 levels as monomeric pT217 declined (**Figure S6M** and **S6N**). A similar pattern was observed in the pT181 species. At 1.5 months, the PSEN1^M146V/WT^ COs had elevated levels of monomeric pT181 without significant HMW bands (**Figures S6I** and **S6K)**. Over time, however, we observed the monomeric pT181 levels losing its elevated levels in the monomeric pool while HMW pT181 progressively increased at the 3- and 6-month timepoints (**Figures 3H** and **3I**). These results highlight that PSEN1 COs in particular progressively accumulate HMW pTau species. Furthermore, these data suggest the fAD CO model may mechanistically capture the time course of the development of pT217 and pT181 oligomer-like HMW species.

Finally, to further corroborate the extent to which the COs develop oligomeric pTau species, we ran a NATIVE PAGE under reducing conditions at ∼4-month timepoint (**Figure S6O**). Probing for pT181 revealed abundant smeared bands at ∼480-720 kDa apparent MW in both mutant COs, most prominently in PSEN1^M146V/WT^.

### AD COs recapitulate signatures of neurodegeneration

To test the hypothesis that the relative progressive loss of mTau and/or monomeric pTau at later time points could be associated with neuronal loss, we characterized cell death in the COs at the 1.5 month and 3 month timepoints via TUNEL (terminal deoxynucleotidyl transferase dUTP nick end labeling) staining^100^ (**Figures S7A** and **S7B**). We complimented this approach by counting nuclei with condensed chromatin to assess apoptotic cell death (**Figure S7C**).^101,102^ We found the AD CO mutants of both genetic backgrounds to have a significantly elevated number of apoptotic cells compared to their isogenic WT COs by the 3 month-timepoint. Importantly, both backgrounds of WT isogenic COs showed consistent and stable numbers: they exhibited neither significant increases in cell death between timepoints, nor any differences when compared to each other at either timepoint (**Figure S7C, *right***). This consistency demonstrates our results are robust and independent of the COs’ genetic background. This finding suggests cells in the mutant AD COs may either have developed increased vulnerability during that time or other emergent mechanisms have led to increased cell death within the mutant COs only.

### AD COs have an abnormally high basal level of autophagosomes

A decrease in autophagic flux has linked to AD pathophysiology, and activation of autophagy has been suggested as a potential therapeutic target for neurodegenerative diseases like AD.^45,47–49^ We therefore characterized basal autophagic flux state in the COs. We performed live imaging utilizing commercially available DAPRed (Dojindo).^103^ This amphiphilic dye integrates into the double membrane of autophagosomes. DAPRed fluoresces upon hydrophobic membrane incorporation, labeling autophagosomes and autolysosomes. We compared the number of DAPRed puncta in the vehicle control (DMSO) relative to bafilomycin A1 (BafA1) exposure, which prevents autophagosomes from fusing with lysosomes to create an acidic degradation compartment in the autolysosome. The BafA1 condition not only ensures the endo-lysosomal flux is not blocked—as otherwise there would be no difference in the number of autophagosomes accumulating after BafA1 exposure—but it can also estimate the number of autophagosomes processed through the lysosomal pathway. We found an increased number of autophagosomes converted to autolysosomes in all the AD mutants (**Figure 4A** and **4B**; cf. difference between +BafA1 across genotypes). Despite having more autophagosome load, the lysosomal flux system was not clogged or impaired, as demonstrated by the BafA1 exposure significantly increasing the number of autophagosomes (**Figure 4A** and **4B**; cf. difference between DMSO and BafA1 condition). This suggests an increase in basal macroautophagic flux in fAD, likely compensating for aberrant proteostasis early in the course of disease, as studied in our CO model system.

**Figure 4:**
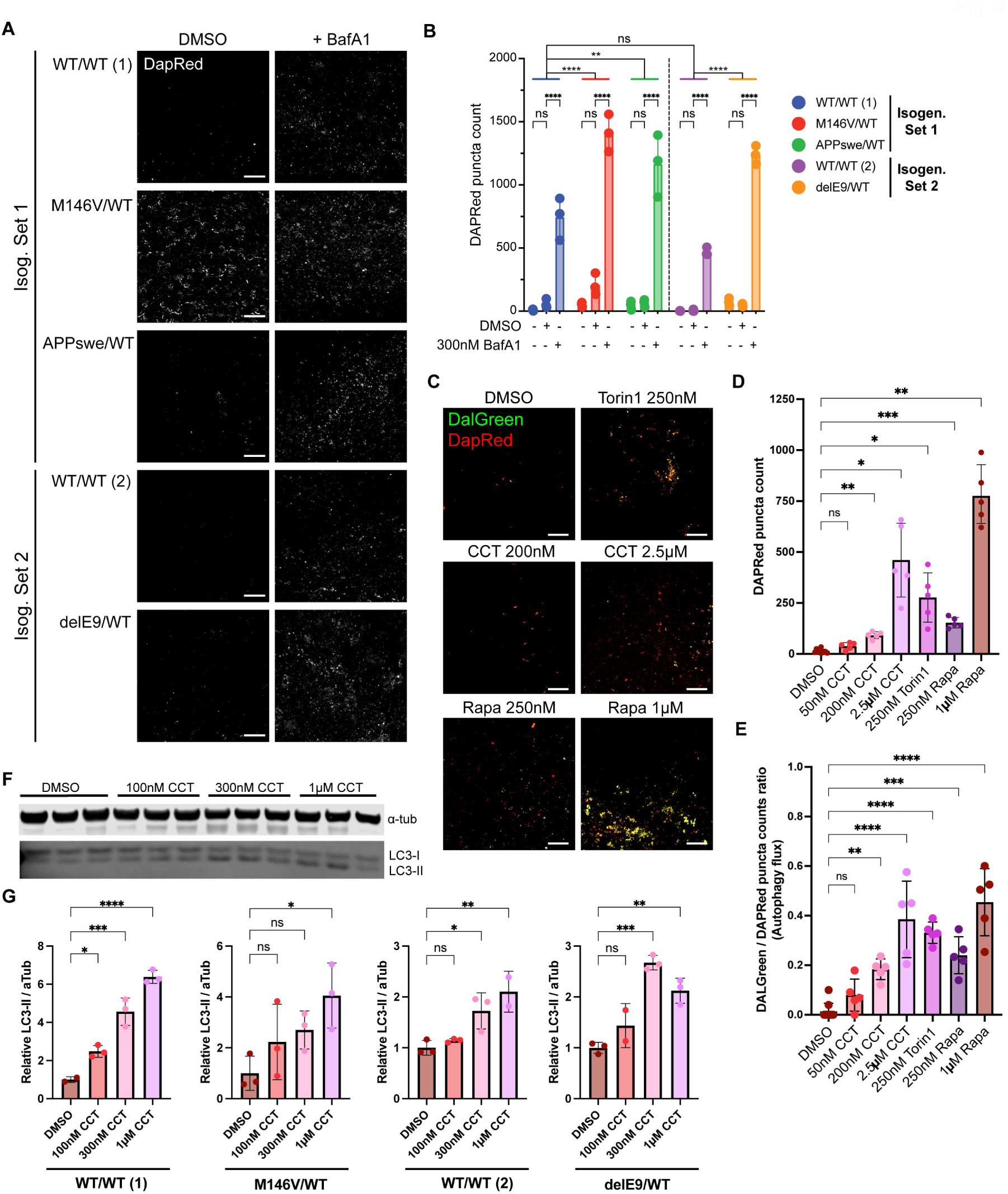
Autophagy can be efficiently activated in the AD COs. (A and B) Autophagosome staining via DAPRed live staining of COs in Neural Media (N.Media), with an overnight treatment of DMSO or 300 nM of Bafilomycin A1 (BafA1) as controls, along puncta quantification. Scale bar, 50 µm. (B) Quantification of average DAPRed+ puncta count per image. n = 3 COs per condition. Analysis by 2-way ANOVA with Dunnett’s and Sidak’s post-hoc tests for inter- and intra-genotype comparisons, respectively. (C) Representative immunofluorescence (IF) staining images of DALGreen and DAPRed signal in COs (pictured here: PSEN1^delE9/WT^) after a 24-hour treatment with either different concentrations of autophagy activators Rapamycin (Rapa), CCT020312 (CCT), Torin 1 (Torin1), or vehicle. Scale bar, 50 µm. (D and E) Quantification of the average IF DAPRed+ puncta count per image (D) and DALGreen to DAPRed puncta counts ratio (E). n = 5-10 images from 3-4 COs per condition from 1-2 independent experiments. Representative WB of the dose-dependent LC3-II abundance increase in COs (pictured here: WT/WT (2)) after a 24-hour treatment with CCT. (G) WB Quantification of the relative LC3-II abundance in 24-hour-treated COs, normalized to their respective DMSO values. Data are mean ± SD. Analyses by ANOVA with Dunnett’s post-hoc test.

### Increases in AD CO disease-correlated pTau are reversed by the novel lysosomal flux activator, CCT020312

Given the observed elevated autophagosome load in the fAD COs, we posited that pharmacological activation of lysosomal flux could enhance macroautophagy efficiency, thereby reducing the levels of pathogenic protein aggregates associated with the disease. Recently, compound CCT020312 (denoted CCT hereafter) was identified from a cell-based high-content screen and validated as an mTOR inhibitor-independent lysosomal flux activator.^104^ This was of particular interest to us given mTOR inhibition’s unwanted side effects such as immunosuppression. CCT was previously shown to dose-dependently reduce insoluble pTau levels in a hiPSC-derived 2D neuronal model of Tauopathy and decrease Aβ peptides in direct-differentiated neurons.^104^ The vulnerability of the 2D neuronal cultures of Tau-P301L neurons to stressors like rotenone and Aβ also exhibited substantially less cytotoxicity with CCT pre-treatment. Thus, we hypothesized that autophagy activators like rapamycin—an mTOR inhibitor^47^—and CCT could potentially reverse the clinically relevant phenotypes of pTau accumulation that we identified in our fAD COs.

Next, we assessed the ability of known mTOR-dependent autophagy activators, like Torin 1 and rapamycin (Rapa), or the mTOR-independent drug CCT to increase lysosomal flux in the AD COs at comparable concentrations to other in vitro models.^105^ To rigorously assess lysosomal flux, we made use of DAPRed and DALGreen (Dojindo) in conjunction. While both DAPRed and DALGreen integrate into autophagosomes, the fluorescence of DALGreen is greatly enhanced in acidic environments, thus labeling autolysosomes where the pH is lower. This enables the quantification of the relative ratio of mature autolysosomes and early autophagosomes; that is, live assessment of flux from autophagosome formation to post-fusion autolysosome maturation. We observed a significant dose-dependent increase in both autophagosomes (DAPRed puncta) and autophagic flux (DALGreen/DAPRed punctae ratio) after 24-h treatment with Torin 1, Rapa, or CCT (**Figures 4C-E**). Our assessments agreed that 250 nM Torin 1, and doses as low as 250 nM Rapa or 200 nM CCT were sufficient to significantly increase autophagic flux in the COs after 24 hours.

We further validated the effect of these autophagy activators via LC3-II quantification through western blotting (**Figures 4F**). We found that, in general, concentrations as low as 100 nM CCT led to increased LC3-II accumulation in the WT/WT (1) COs, but there was some variability depending on the CO genotype. (**Figure 4G**). Overall, the live dye and LC3-II assessments suggest concentrations of ∼300 to 1 µM CCT are required to robustly activate autophagy in the COs regardless of the genotype.

Next, we set out to study the therapeutic effects of chronically treating the COs with autophagy activators, employing various dosing regimens. We began with mTOR-dependent Rapa using two different treatment regimens of 2 to 4 weeks based on prior similar treatments on a different hiPSC-derived 2D neuronal model.^104^ Specifically, we tested dosing twice with 250 nM Rapa at day 0 and day 13 over the span of 2 weeks, as well as a more frequent lower dose regimen of 125 nM Rapa every 4 days over the span of 4 weeks, after which we lysed the COs to assess the effect on pTau (**Figures S8A** and **S8B**). We found that both regimens yielded nearly identical results and thus were combined for analysis. The Rapa treatment increased levels of mTau in the PSEN1^M146V/WT^ COs (**Figure S8C**), while significantly decreasing levels of HMW pT217 in all COs (**Figure S8D**). Rapa also decreased pT181 in the APP^Swe/WT^ COs (**Figure S8E**). Overall, we found that rapamycin was effective at reducing pTau compared to vehicle.

Similarly, we aimed to determine whether chronic treatment with the mTOR inhibition-independent autophagy activator, CCT, could replicate the decrease in tau pathology observed with rapamycin treatment. This comparison was particularly important because inhibition of mTOR has significant side effects that may not be tolerated well clinically,^47,106^ motivating our development of an mTOR-independent mechanism to activate autophagy. We therefore conducted a series of experiments to comprehensively evaluate the effects of CCT on the COs (**Figure 5A**). For a 4-week-long treatment period, multiple batches of COs starting at 2-month timepoint were dosed every 4 days with CCT or vehicle prepared in fresh neural medium (NM) at every other medium change. As a positive control to complement the rapamycin findings, we used a second mTOR-dependent activator of autophagy, Torin 1, at a concentration of 200 nM.

**Figure 5:**
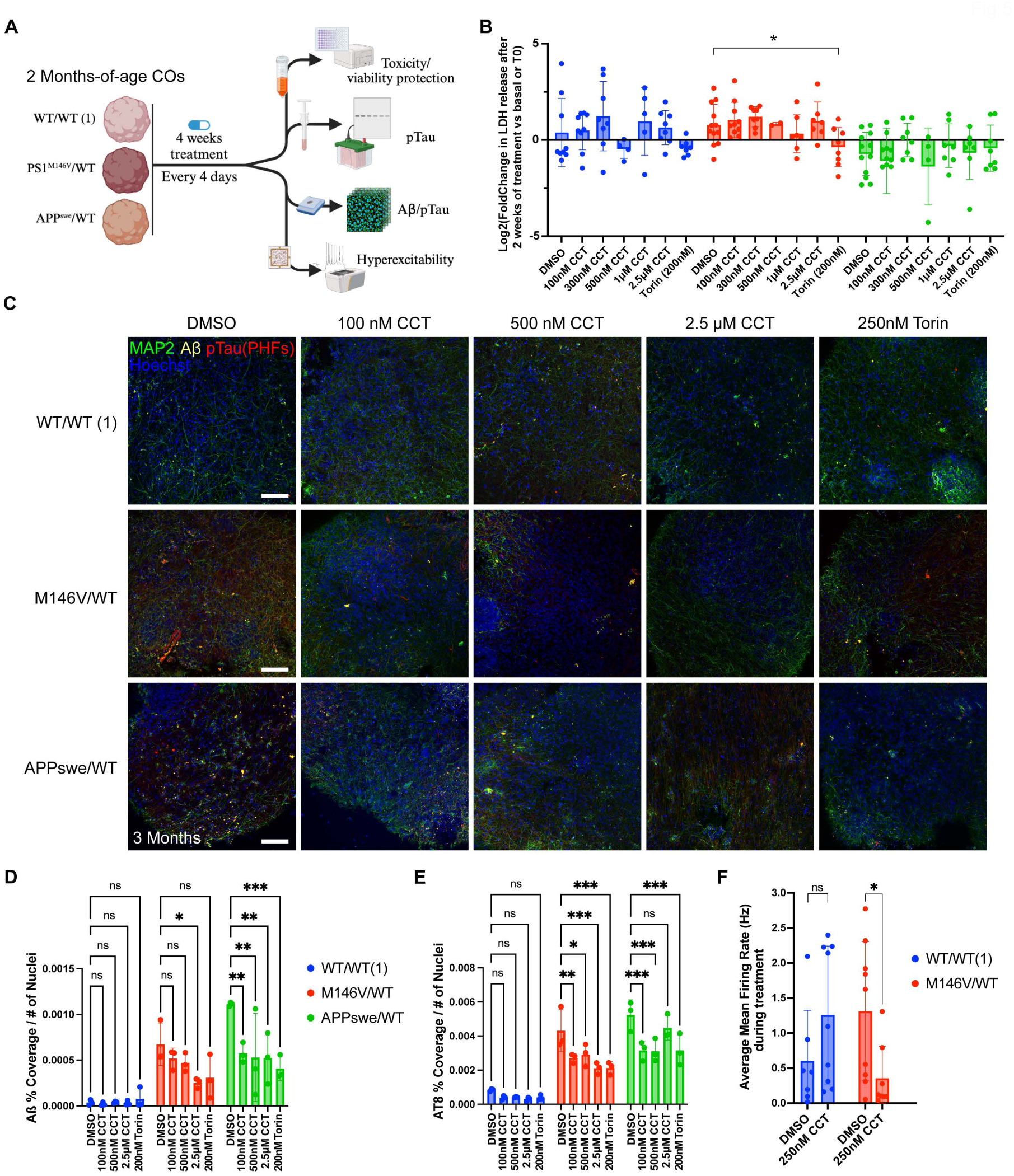
Chronic mTOR-independent autophagy activation reduces AD-associated Aβ and pTau aggregates, and hyperexcitability. (A) Diagram describing the experimental set-up of the 4-week-long treatment of 2 months-of-age COs (3-month timepoint by end of treatment). (B) Measured change in the conditioned media levels of LDH (U/ml) at the end of treatment (4 weeks), compared to their basal levels at treatment day 0 (T0) (LDH at end of treatment minus LDH at T0). Data was Log-transformed to achieve normality. (C) Representative immunofluorescence (IF) images of the COs at the end of the treatment period. Scale bar, 100um. (D) Quantification of the beta amyloid percent coverage normalized to nuclei number of the COs. n = 3 COs per condition. (E) Quantification of the Pre-Helical Filament aggregated pTau (AT8^+^) percent coverage normalized to nuclei number of the COs. n = 3 COs per condition. (F) Average weighted mean firing rate of the WT/WT (1) and M146V/WT COs in the 12-24 hours period after treatment. n = 7-9 COs per genotype from 2 independent experiments. Data are mean ± SD. Analyses by ANOVA with Dunnett’s post-hoc test (B, D, and E) or Kruskal-Wallis 2-tailed non-parametric test (F). See also Figures S9, S10, and S11.

Potential toxicity due to the long-term dosing of the drugs was tracked by measuring LDH release in conditioned media from pre-treatment day zero (T0) to the mid-, and endpoint every two weeks. We did not find any dose-dependent toxicity in any of the CO genotypes, although, notably, we observed relatively lower levels of LDH released in PSEN1^M146V/WT^ COs treated with 200 nM Torin 1 (**Figure 5B**). Next, we assessed immunohistologically the COs for Ab and PHF pTau pathology (**Figure 5C**). We not only confirmed once again elevated levels of Ab and AT8+ signal in both PSEN1^M146V/WT^ and APP^Swe/WT^ COs compared to WT, but we also found a clear, dose-dependent decrease in both Ab (**Figure 5D**) and AT8+ PHF signal with CCT treatment (**Figure 5E**). This demonstrates the ability of both CCT and Torin 1 (used here as a positive control) to directly reduce AD pathology.

Previously, CCT has been regarded as an activator of the PERK arm of the Unfolded Protein Response (UPR).^107–112^ Recent work, however, from our groups has shown that CCT enhances lysosomal flux at lower concentrations than those required to activate PERK.^104^ To verify this was the case in here in our human CO model, we treated for both WT/WT and PSEN1^M146V/WT^ mutant COs for one week with CCT at concentrations ranging from 100 nM to 25 µM. COs were then lysed and probed for Activating Transcription Factor 4 (ATF4), a canonical downstream target of PERK/UPR activation.^113,114^ We found that CCT did not have any effect on ATF4 translation below 10 µM (**Figure S9A**). As a postive control, 500 nM Thapsigargin, a well-validated UPR activator,^115,116^ induced a consistent increase in ATF4 signal in both CO genotypes. Critically, pretreatment or co-treatment of CCT with Integrated Stress Response Inhibitor (ISRIB), a small molecule acting downstream of PERK activation and a potent counteractor of its effects,^117^ had no effect on the ability of CCT to reduce PHF load in AD COs (**Figure S9B**). Thus, we conclude the therapeutic effects of CCT observed here at concentrations ≤2.5 µM are independent of its ability to activate PERK, which is only seen at higher concentrations.

Following up on our aforementioned finding of pTau accumulation in the PSEN1^M146V/WT^ mutant, 4-week CCT-treated CO lysates were also biochemically assessed for pTau changes (**Figure S10A**). The results showed that the mTau levels were not significantly affected by lysosomal flux activator treatment in either genotype at this timepoint (**Figure S10B**). In contrast, PSEN1^M146V/WT^ COs manifest a significant, dose-dependent reduction in pT181 after treatment with 1 – 2.5 µM CCT (**Figure S10C**). Torin 1 (250 nM) appeared to have a similar albeit somewhat less effect, but, unlike CCT, is also a mTOR inhibitor at this or higher concentrations, which would produce substantial clinical side effects. Finally, CCT had no statistically significant effect on pT217 levels (100 kDa band). Consistent with our findings that pT217 at this MW only accumulates in the PSEN1^M146V/WT^ COs at later timepoints, these results are consistent with the notion that the mTOR-independent lysosomal flux activation by CCT may preferentially target abnormally elevated pTau species.

### Chronic treatment with lysosomal flux activator, CCT, rescues mTau, monomeric pTau, and synapses in PSEN1^M146V/WT^ COs

We identified between the 6-month and 9-month timepoints a period during which the PSEN1^M146V/WT^ AD COs appear to lose synaptic density, as evidenced by our biochemical assessment of presynaptic marker Synapsin I (**Figure 6A**) and consistent with our prior electrophysiological and histological assessments on similar AD COs.^43,44^ This observation, combined with assessments of cell death and mTau reduction, suggested the PSEN1^M146V/WT^ mutant COs underwent progressive synaptic loss typical of neurodegeneration compared to isogenic WT and to a greater degree than the APP^swe/WT^ COs. To assess whether CCT could affect this pathological progression, we conducted a 6-week treatment regimen initiated in COs at 6 months of age and ending at the 7.5-month timepoint (**Figure 6B**). To assess viability, we measured LDH levels in conditioned media at the pre-treatment, 1-week, 2-week, and 4-week timepoints (**Figure 6C**). Although all conditions showed some increase in LDH release after the first week, the PSEN1^M146V/WT^ COs manifested a significantly higher increase in LDH released, suggesting their heightened vulnerability. Notably, CCT treatment did not exhibit toxicity compared to DMSO controls at any timepoint. On the contrary, we observed a significant reduction in LDH release relative to pre-treatment levels in the PSEN1^M146V/WT^ COs treated with 100 nM CCT as soon as one week after treatment (**Figure 6C**, ***right***).

**Figure 6:**
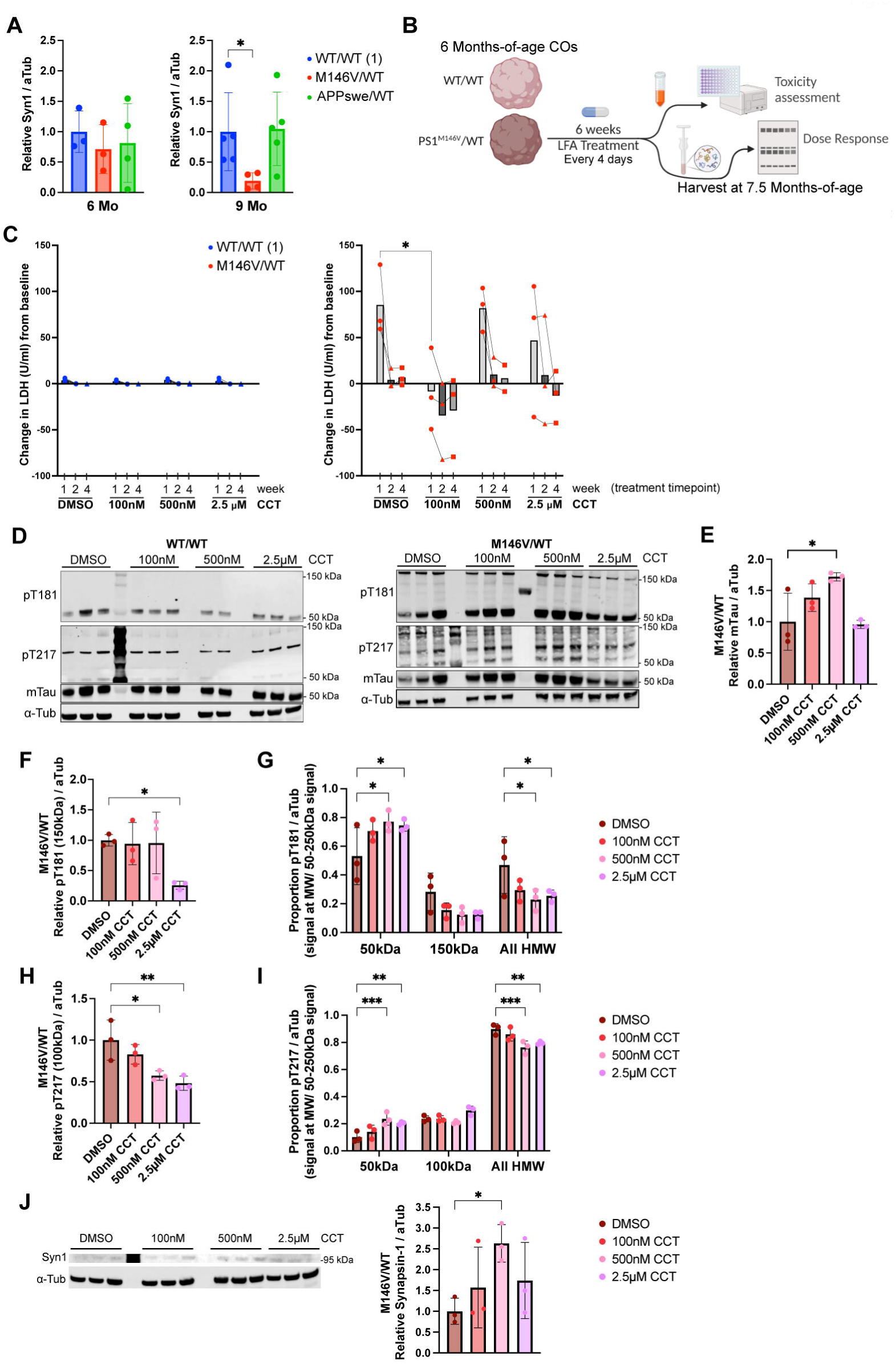
Chronic mTOR-independent autophagy activation with CCT rescues mTau, monomeric pTau, and synapses in the M146V mutant. (A) Immunoblot quantification of synaptophysin-1 (Syn1 normalized to a-Tub) in CO lysates at 6 and 9 months-of-age. n = 3-4 COs per genotype. For each timepoint, Syn1 levels shown relative to WT/WT (isogenic set 1) levels. Unpaired t test with Welch’s correction with post-hoc FDR-adjusted p-values: p=0.0461 (M146V-vs-WT(1)), p=0.9082 (APPswe-vs-WT(1)). (B) Schematic describing the treatment of 6 months-of-age COs with CCT and its assessment at the end of a 6-week treatment regimen (harvested at 7.5 month-timepoint). (C) Quantification of LDH in conditioned media after 1, 2, and 4 weeks of treatment with DMSO vehicle vs. 100 nM, 500 nM, or 2.5 µM CCT for WT/WT (1) or M146V/WT. n = 5-6 COs per condition for WT/WT (1); n = 3 COs per condition for M146V/WT. (D) Representative immunoblots of WT/WT and M146V/WT CO lysates for pT217, pT181, mTau after 6-week treatment. (E) Quantification of monomeric Tau (mTau normalized to a-Tub) in CO lysates after 6-week treatment relative to DMSO. (F) Quantification of phosphorylated Tau T181 (pT181 normalized to a-Tub) in CO lysates after 6-week treatment relative to DMSO. (G) Quantification of pT181 bands (normalized to a-Tub) at various molecular weights (MW); specifically, at the monomeric band (50 kDa), major validated high-molecular-weight (HMW) band (150 kDa), and all HMW signals (75-250 kDa). (H) Quantification of phosphorylated Tau T217 (pT217, normalized to -Tubulin) relative to DMSO levels. (I) Quantification of pT217 bands (normalized to a-Tub) at various molecular weights (MW); specifically, at the monomeric band (50 kDa), major HMW band (100 kDa), and all HMW signals (75-250 kDa). (J) Representative immunoblot of DMSO vehicle-vs. CCT-treated M146V/WT CO lysates for Syn1 and a-Tub after 6-week treatment (*left*) with quantification (*right*). n = 3 COs per condition. Data are mean ± SD. Analyses by ANOVA with Dunnett’s post-hoc test.

When monitoring pTau, we noted that the HMW bands of pTau became significantly more complex and developed smearing in the PSEN1^M146V/WT^ CO lysates (**Figure 6D, *right***), resembling more closely the signature of oligomeric pTau pathology in human AD patient brains.^96^ Surprisingly, CCT dose-dependently increased the abundance of mTau, reaching significance at the 500 nM dose (**Figure 6E**). This is important given our finding of mTau dramatically decreasing in the PSEN1^M146V/WT^ COs beyond the 4.5-month timepoint in the absence of treatment. Furthermore, CCT dose-dependently decreased the major HMW pT181 and pT217 bands (**Figures 6F-I**). Consistent with our hypothesis of CCT-activated autophagy targeting abnormally elevated pTau, 500 nM and 2.5 µM CCT treatment not only decreased the abundance of the major HMW pT181 and pT217 bands (**Figures 6F and 6H**), but also simultaneously shifted the relative proportion of HMW-to-monomeric pTau species, as demonstrated by the increase in monomeric pT217 and pT181 bands (**Figures 6G and 6I**). This finding is consistent with the notion that CCT-activated autophagy may digest SDS-insoluble oligomers into SDS-soluble monomers.

Finally, in PSEN1^M146V/WT^ COs we found that 500 nM CCT significantly increased the levels of Synapsin I compared to DMSO vehicle (**Figure 6J**), suggesting a beneficial effect on synapse number. Taken together, these results nominate CCT as a potential therapeutic worthy of further testing in a human disease context, at least for PSEN1^M146V/WT^ mutations.

### Pathologic neuronal hyperexcitability in the PSEN1^M146V/WT^ AD COs is corrected by the lysosomal flux activator, CCT

Hyperexcitability has been identified as an early and important contributor to the pathogenesis of AD, affecting Tau aggregation and synapse loss.^43,74,118–120^ Using a microelectrode array (MEA) recording system (Axion Bioystems), we attached individual COs to single MEA wells to enable real-time readings of their basal electrical activity. By resistivity measurements, we verified that CO attachment to the MEA plate was not different between mutants and WT. By the 3-month timepoint, the PSEN1 CO mutants produced abnormally high levels of action potential firing and bursting, as evidence of hyperactivity (**Figure S11A**). This was in line with results we had previously reported on hypersynchronous activity found in PSEN1^M146V/WT^ mutant COs compared to isogenic WT COs,^43,44^ so we moved directly into assessing whether CCT could also correct hyperexcitability in the AD COs.

Treating every 6 days for a month, we assessed viability and weighted mean firing rate (**Figure S11B**) with an extended chronic dosing regimen to assess long-term the viability of the COs. Based on our previous treatment successes, we compared dosing the isogenic PSEN1^M146V/WT^ COs and their isogenic WT COs with vehicle DMSO (**Figure S11C**) vs. 200 and 300 nM doses (∼250 nM, hereafter) CCT (**Figure S11D**). We found a genotype-specific decrease in the PSEN1^M146V/WT^ CO activity compared to isogenic WT control. While WT COs were not affected, the 250 nM CCT dose significantly reduced PSEN1^M146V/WT^ CO hyperactivity during the treatment period (defined as the stable activity period between 12-24 hours after a media change with the drug/vehicle). The activity remained relatively lower than basal levels during the first few days following drug wash out (24-72 hr post drug). Electrode impedance measurements, reflecting both attachment and membrane integrity within the COs, showed surprising results. During the 24-72 hr post drug period, DMSO (vehicle alone)-treated PSEN1^M146V/WT^ COs manifested a significant drop in impedance compared to their basal levels (**Figure S11E**). This did not occur in the DMSO-treated WT COs; the effect occurring only in the PSEN1^M146V/WT^ COs again reflects the increased vulnerability of AD COs. Notably, 250 nM CCT treatment rescued this effect on the AD COs (**Figure S11F**). This finding supports our interpretation that the decrease in hyperexcitability observed with CCT treatment in the PSEN1^M146V/WT^ COs was not due to a decrease in cell viability. Taken together, the results are consistent with the notion that CCT limited the AD-related hyperexcitability and viability loss phenotypes. Moreover, our findings suggest that the CO platform is a viable model for assessing the effects of autophagy activators like CCT in a human context in real time using relevant electrophysiological as well as biochemical readouts.

## DISCUSSION

In this study, we have leveraged hiPSC-derived COs to create a scalable and consistent in vitro human model for fAD vs. isogenic controls, using it to focus on the earliest AD-related pathology. We demonstrate this model succeeds in capturing both the hallmark increase in Ab and phosphorylated Tau proteins with synaptic marker loss and cellular death, as well as the recapitulation of pathophysiologic neuronal hyperactivity observed in pre-symptomatic PSEN1 mutant and APOE4 variant carriers via hippocampal fMRI, and early in human AD brains by electroencephalogram (EEG).^5,8, 121,122^

The AD COs developed here not only replicate the early molecular, transcriptomic, and synaptic abnormalities previously reported in clinical human AD patients,^2,73,89,96,123,124^ but also reveal distinct molecular signatures depending on the familial AD mutations present. Specifically, we show the PSEN1 mutant COs developing the strongest phenotypes in most of our characterization, including elevated early levels and progressive accumulation of HMW pT217 and pT181, synaptic loss, and neuronal hyperexcitability. This suggests that mutations in the PSEN1 and APP genes drive unique aspects of AD pathology, which may help explain their differential associated clinical prognosis. Indeed, the PSEN1 mutants were the only ones that had an elevated Ab_1-42_,/ Ab_1-40_ ratio, which is considered a stronger prognosis diagnostic than just elevated Ab_1-42_ alone, which APP^swe/WT^ COs had.^125^ PSEN1 mutations are also associated with significantly more aggressive early onset dementia with age of onset of 24-55 years of age,^37,38,123^ while the APP^Swe/WT^ mutation is associated with slower age of onset of 40-65 years of age.^123^ Additionally, NFL, a marker for axonal damage has been found to increase earlier in PSEN1 carriers compared to APPswe carriers.^82^ The fact that the Ab_1-42_/ Ab_1-40_ ratio also followed over time the same mutant-specific progressive decline as reported in CSF clinical studies decades before symptom onset,^80–82^ suggests the CO model to be reproducing real progressive processes at an accelerated pace.

The relative decrease in monomeric phosphorylated Tau was an intriguing finding in our study. Clinical studies on AD patients have found significant reduction in the abundance of APP and Tau protein in CSF microvesicles.^125^ We hypothesize the COs may be reproducing a process early in the disease etiology, by which specific pTau may be irreversibly oligomerizing, leading to its decrease in SDS-soluble extraction. The formation of phosphorylated Tau dimer and higher order intermediate oligomers as they form Tau fibrils has been previously explored.^126–128^ New evidence suggests these low-order oligomers may be the most hazardous form of Tau oligomers.^129,130^ The field in general agreed that hiPSC-derived neurons do not exhibit Tau multimerization.^128,131^ To our knowledge, we are the first to report validated low-order, reduction- and SDS-resistant pTau oligomers in any *in vitro* model.

Recently, AD patient-derived soluble HMW tau was found to drive abnormal neuronal bursting in mice hippocampal neurons.^132^ The fact the COs exhibit both of these suggests they may be an accessible human model to study electrophysiologically-relevant pathology lacking in other models. Additionally, our findings reveal starkly different effects in excitatory neurons compared to inhibitory ones, which may explain why patients develop excitatory-inhibitory imbalance during the disease and highlight potential new therapeutic targets in the transcriptomic signatures of these neuronal subtypes. Recent studies support this notion.^133–136^

A key limitation of this study is the inherent immaturity of hiPSC-derived cerebral organoids (COs), which despite showing senescent signatures, cannot fully recapitulate the epigenetic age of adult brain cells. The absence of microglial cells further restricts the model’s ability to reproduce inflammatory disease processes critical for protein aggregate clearance and disease progression, potentially limiting our findings to pre-inflammatory early AD stages. The organoid models have been previously suggested to involve non-physiologic stressors,^26^ and the accelerated development of AD pathology in COs compared to human brain supports this notion, but these characteristics may actually be advantageous. The presence of cellular stressors-including the ferroptosis and senescence-related stress we identified (elevated p16, p21, and SASP genes)^71,72,137^ may effectively mimic age-related processes that drive AD progression.^26,138^

We posit that the presence of cellular stressors and senescence signatures could contribute to accelerating AD-relevant processes in the COs, despite the potential presence of unrelated non-physiologic stressors.^17^ Furthermore, the three-dimensional structure of organoids provides a more physiologically relevant microenvironment in a human context that supports the development of AD-related pathologies, such as the Tau hyperphosphorylation and aggregation we observe, resembling the high-molecular-weight oligomers observed in human postmortem AD brain. This may explain the emergence of more complex functional abnormalities, such as the differential effect in interneurons, hyperexcitability, and cell death.

Our study demonstrates that chronic treatment with lysosomal flux/autophagy activators—including mTOR-dependent agents like rapamycin and Torin 1, but also mTOR-independent drugs like CCT020312—can reduce AD-like phenotypes, including the accumulation of Ab and pTau over a two to six-week treatment period. We show that CCT activates autophagy without activating PERK or the UPR, and that this activity reduces aberrant HMW pTau species. Critically, autophagy activators not only reduced pathogenic protein burden but also preserved cellular viability and electrophysiological properties of the AD COs. Despite chronic, weeks-long exposure to these therapeutic agents, COs did not show toxicity; in fact, treatment rescued synapses and prevented hyperexcitability and cell death. Correlated with this neuroprotective effect of CCT, we observed an increase in Tau monomers and a concomitant decrease in HMW pTau species, suggesting that the oligomers may drive pathologic toxicity rather than total pTau load. Given the growing evidence that autophagy activation can also reduce neuronal senescence and its potential involvement in neurodegeneration,^139^ these findings strengthen the evidence for autophagy modulation as a promising therapeutic avenue for AD and provide insights for dosing strategies in future clinical applications.^45,47,48,50,106,140,141^

Finally, we showcase the use of real-time MEA recordings in COs as a useful platform for preclinical drug screening and investigation of the potential therapeutic effectiveness of drugs. The generation of hiPSC-derived COs from various additional AD-related mutations or sporadic cases should also facilitate personalized medicine approaches to AD treatment.

## STAR METHODS

### RESOURCE AVAILABILITY

#### Lead Contact

Any additional information and/or requests should be directed to the lead contacts, or.

#### Materials availability

The unique materials generated in this study are available from the lead contact upon completing a Materials Transfer Agreement.

#### Data and code availability

Single-cell RNA-seq data is deposited at GEO, accession number GSE301700.

No significant custom code was generated in this study, though the specific basic R scripts used are available upon request. Mass spectrometry data will be available upon request. Any additional information required to use or reanalyze the data reported in this study is available from the lead contact upon request.

### EXPERIMENTAL MODEL AND SUBJECT DETAILS

#### Human iPSC Lines

The use of human cells had the approval of the institutional review board associated with the Scripps Research Institute. We used the previously established and characterized^1, 2^ isogenic heterozygous PSEN1^M146V/WT^ and APP^Swe/WT^ mutant hiPSCs, along with their WT 7889SA parental line (M146V/WT, APPswe/WT, and WT/WT (isogenic set 1)), obtained from the Tessier-Lavigne laboratory (for mutant lines) and the New York Stem Cell Foundation Research Institute (for the WT/WT line). For an additional genetic background, we used the previously established^3, 4^ PSEN1^ΔE9/WT^ heterozygous hiPSC and its WT parental line (delE9/WT and WT/WT (isogenic set 2)), originally obtained from the Goldstein laboratory at the University of California, San Diego. Both genetic backgrounds were of male sex. Pluripotency was also confirmed via immunolabeling with pluripotent cell markers NANOG, OCT4, and TRA (data not shown). Euploidy was confirmed via both G-banding (WiCell) and qPCR (StemCell Technologies Kit), which showed no significant gain of karyotypic mosaicism in the hiPSCs used in this study.

### METHOD DETAILS

#### hiPSC Maintenance and Quality Controls

The hiPSC sets were maintained in parallel on vitronectin-coated (Thermo Fisher) plates with the culture medium mTeSR plus (StemCell Technologies) being replaced every 24 hours. The hiPSCs were carefully cultivated, morphologically abnormal cells were manually removed when needed, and cells were passaged via non-enzymatic colony dissociation with ReleaSR (Thermo Fisher) every 5-6 days. The cells were routinely checked for mycoplasma and maintained for no more than 13 passages at a time.

#### Cerebrocortical Organoid (CO) Culture

To make the cerebral organoids (CO), an optimized protocol based on Sergiu Paşca’s method was developed.^5^ hiPSCs between 75 and 90% confluence were first dissociated with Accutase into a single-cell suspension and seeded into an Aggrewell plate at a 4 million cells per well density in mTeSR plus media supplemented with a *CEPT* cocktail^6–8^ (50 nM Chroman I, 5uM Emricasan, 1x Polyamine supplement, and 0.7 µM Trans-ISRIB) to form embryoid bodies (EBs) for 24 hours in an 5% CO_2_ incubator at 37 °C. EBs were then transferred to ultra-low attachment 6-well plates (Corning) shaking in an orbital shaker at 60 RPM. They were maintained in a 50/50 mixture of Gibco’s Essential 6 medium and hESC medium (DMEM-F12, 20% Knockout Serum Replacement, 1% Glutamax, 1% MEM-Non-essential Aminoacids, and 0.1 mM 2-mercaptoethanol)^9, 10^, supplemented with 2.5 μM Dorsomorphin, 10 μM SB-431542, and 2.5 μM XAV-939 for 6 days. The medium was then replaced with Neural Media-A supplemented with 20 ng/ml of EGF_2_ and 20 ng/ml FGF_2_. From day 25 to 43, the organoids underwent maturation through replacing the supplements with 20 ng/ml BDNF and 20 ng/ml NT-3; after which they were maintained in Neural media-A until harvest. Importantly, around day 35 the organoids were transferred to ultra-low attachment 10 cm dishes and kept shaking at an orbital shaker tilted at a 3.5° angle. The COs were monitored and split as needed to maintain consistent media consumption/acidification. CO cultures were tested for mycoplasma every 1-2 months.

#### CO Brightfield Imaging

Transmitted light images of live representative COs were taken via an EVOS M5000 Microscope (Invitrogen) with a 2x or 4x objective and phase ring on the brightfield setting. Light intensity/exposure was adjusted to maximize the images’ contrast each imaging day independently but maintained constant across conditions within each day. CO size (area) quantification was done via Fiji with the Cell Magic Wand plugin.

#### Immunohistology of COs

Cerebrocortical organoids were washed 2x in PBS and fixed in a cold solution of 4% PFA in PBS overnight at 4 °C. Organoids were then washed in a 300 mM Glycine solution to neutralize the remaining PFA and washed 2x in PBS for 15 minutes each before being dehydrated through progressive equilibration in 15% and 30% sucrose in PBS overnight. Organoids were then embedded in tissue-freezing medium (General Data) in a cryo-mold and flash frozen in liquid nitrogen–cooled isopentane and stored at -80 °C until processing. Fluorescence immunohistochemistry was performed on 20 µm cryostat-sliced samples using standard protocols. Briefly, blocking and permeabilization were done with 3% BSA + 0.3% Tx-100 for 10 minutes at RT, after which samples were incubated overnight at 4 °C with primary antibodies diluted in the same blocking buffer (dilutions in Key Resources Table). Then, samples were washed with PBS and incubated with Alexa Fluor-labeled secondary antibodies and Hoechst nuclei stain. Finally, samples were washed three times in PBS, coverslipped with Dako Fluorescence Mounting Medium, and sealed. Secondary antibody-only controls were carried out as negative controls for each experiment. A Nikon C2 confocal microscope was used to perform the imaging, acquiring at least 3 Z-stack captures in randomly selected fields for each replicate condition.

#### Western Blot Characterization

Cerebral organoids of 1.5-, 3-, and 6-month timepoints were washed 3x in PBS and then homogenized in protease and phosphatase inhibitor cocktail-supplemented RIPA or 2% SDS lysis buffer (2% SDS in 150mM Tris-HCl, 7.6 pH). Results did not vary significantly based on the choice of lysis buffer. To ensure a complete protein extraction, after homogenizing the COs, the samples were directly pulse ultrasonicated (∼24 seconds) before centrifuging at 20,600 x g for 20-30 minutes. Supernatant was saved into a new tube and protein was quantified and normalized using Pierce BCA assay (Thermo Fisher). Next, SDS-PAGE was conducted –using SDS-free Laemmli sample buffer– with Bolt Bis-Tris 4-12% gradient polyacrylamide gels (Thermo Fisher). Gel was washed in 20% EtOH for 20 minutes, washed in 10% MeOH Transfer Buffer for 10 minutes, and then dry transferred onto PVDF membranes using an iBlot 3 Western Blot Transfer Device (Thermo Fisher). The membrane was then washed with water and stained with Ponceau S stain as protein loading and transfer control. Then, the membrane proceeded with washing with TBST, blocking with Li-Cor TBS blocking buffer, and incubating overnight at 4°C with primary antibodies diluted in the blocking buffer with 0.02% TWEEN-20 (antibody dilutions in Key Resources Table). Membranes were then washed 3x with TBST prior to incubating with secondary antibodies (1:10K) for 1-2 hours at RT. After washing 3x with TBST, membranes were scanned in an Odyssey CLx scanner (LI-COR). After washes and prior to scanning, membranes were stored in TBS (BioPioneer Inc.). Densitometry of protein bands was performed in ImageStudio.

#### High-Molecular-Weight phospho-Tau bands preparation for mass spectrometry (MS) validation

Fresh ∼3 months-of-age CO lysates were prepared as above, homogenized in 1% NP-40 buffer with protease and phosphatase inhibitors. Enrichment was done via incubating the lysate supernatant overnight with protein G Dynabeads (Invitrogen) pre-loaded with fresh antibody, following manufacturer recommendations. After determining protein concentrations on the clear supernatant, 2 µg of protein were loaded for SDS-PAGE in technical duplicates. Duplicate lanes were cut and stained with Coomassie blue. Using a scalpel, the bands were cut at the relative molecular weights where phospho-Tau pT181 and pT217 had been previously observed (∼105 and ∼155 kDa). These samples were then in-gel digested and prepared for MS analysis. Gel with remaining uncut samples was processed through normal WB for pT217 and pT181 to ensure pTau+ bands were successfully cut out.

#### In-gel digestion sample preparation

In-gel digestion was performed following a previously published protocol with minor modifications. Briefly, gel bands corresponding to approximately 55 kDa, 105 kDa, and 150 kDa, as identified by western blot, were excised and cut into 1 mm³ cubes. Cysteine reduction and alkylation were performed by incubating the gel pieces in 5 mM Tris(2-carboxyethyl)phosphine (TCEP) at room temperature for 30 min and 10 mM iodoacetamide in the dark for 30 min, respectively. The proteins were digested with MS-grade trypsin/Lys-C protease mix (Pierce) at 37 °C for 18 h with shaking. Peptides were eluted by shaking in buffer containing 50 % acetonitrile in 0.1% trifluoroacetic acid (TFA). Peptides were then desalted using C18 spin columns (Pierce) and dried using SpeedVac.

#### Liquid chromatography and mass spectrometry (MS) analysis

Peptides were resuspended in 20 μl of 0.1% formic acid. Samples were analyzed using an Orbitrap Exploris 480 mass spectrometer coupled to a Vanquish Neo UHPLC system (Thermo Scientific). Peptides were first injected onto an PepMap™ Neo Trap Cartridge (300 μm X 5 mm, C18, 5 μm, 100 Å) and further separated on an EASY-Spray™ PepMap™ Neo column (75 µm X 500 mm, C18, 2 µm, 100 Å) at a flow rate of 350 nL/min with an initial solvent composition of 98% buffer A (0.1% formic acid in water) and 2% buffer B (0.1% formic acid in 80% acetonitrile).Peptide eluted with a linear gradient with increasing buffer B from 2% to 20% over 82 min, followed by an increase from 20% to 35% buffer B for 20 min. The column was then washed with 99% buffer B for 8 min. The column was kept at 45 °C at all times.

Mass spectrometry analysis was performed with Data-independent acquisition (DIA) mode. Full MS1 scans were acquired with mass-to-charge ratios ranging from 380 to 980 m/z at a resolution of 120,000 and an automatic gain control (AGC) target of 3 × 10□. Precursor ions were isolated using 7 m/z window with 0.5 m/z overlap, generating 86 MS2 scans within the 380–980 m/z range. MS2 scans were performed at resolution of 30,000, maximum ion injection time of 54 ms, AGC target of 2 × 10□, and normalized collision energy of 28. MS1 data were acquired in profile mode, while MS2 data were acquired in centroid mode.

The DIA-based mass spectrometry data were processed in Spectronaut 19 using library-free (DirectDIA) analyses with the default search parameters.

#### Native PAGE

Fresh CO lysates were homogenized in 1% NP-40 buffer (0.5x Tris Buffer solution, 20 mM HEPES, 0.01% Digitonin, 1% NP-40) supplemented with protease and phosphatase inhibitor cocktails. To ensure a complete protein extraction, after homogenizing the COs, the samples were directly pulse ultrasonicated (∼24 seconds) before centrifuging at 20,600 x g for 30 minutes. Supernatant was saved into a new tube and protein was quantified and normalized using Pierce BCA assay (Thermo Fisher). Next, NATIVE-PAGE was conducted –using NATIVE sample buffer (Invitrogen)– with NativePAGE 4-16% Bis-Tris gradient polyacrylamide gels (Thermo Fisher). Gel was then wet transferred onto PVDF membranes using a Mini Blot Module transfer system (Thermo Fisher) in NuPAGE transfer buffer (Life Technologies). The membrane was then washed with water and stained with Ponceau S stain as protein loading and transfer control. Then, the membrane proceeded with washing with TBST, blocking with Li-Cor TBS blocking buffer, and incubating overnight at 4 °C with primary antibodies diluted in the blocking buffer with 0.02% TWEEN-20 (antibody dilutions in Key Resources Table). Membranes were then washed 3x with TBST prior to incubating with secondary antibodies (1:10K) for 1-2 hours at RT. After washing 3x with TBST, membranes were scanned in an Odyssey CLx scanner (LI-COR). After washes and prior to scanning, membranes were stored in TBS (BioPioneer Inc.).

#### Autophagy Flux Microscopy Assessment

DALGreen and DAPRed (Dojindo) were used following the manufacturer’s protocol. In brief, DAPRed and DALGreen were diluted to working concentration of 0.1 µM and 0.5 µM in Neural Media, respectively. Immediately before the start of a treatment with compound, working solutions of DAPRed and DALGreen were incubated with COs for 90 min. Bafilomycin A1 (BafA1) was used as a control at a 300 nM concentration and was incubated with COs for a 24 hour treatment. Similarly, autophagy modulating compounds (CCT, Rapa, Torin1) were incubated with COs and imaged 24 hours following the start of the treatment. Live imaging was performed using a Zeiss Cell Discover 7 (CD7) after COs transferred to clear, glass-bottom 24 well plates. As temperature and CO2 levels were maintained in the CD7 during imaging, at least 3 z-stacks per CO were captured in randomly selected fields per sample.

#### Beta Amyloid ELISA

1 ml of fresh media per individual CO was conditioned over 24 hours. Then, the levels of Aβ_1-38_, Aβ_1-40_, and Aβ_1-42_ in the resulting conditioned media were measured using a V-Plex Aβ Peptide Panel multiplex kit (Meso Scale Discovery) following manufacturer’s instructions and recorded using an MESO QuickPlex Instrument (Meso Scale Discovery).

#### CO Dissociation for Single cell RNA-Sequencing (scRNA-seq)

Individual COs of ∼7 weeks and 3 months of age were washed 4x in PBS and then dissociated with a Worthington Papain Dissociation kit (Worthington Biochemical Corp.) following a modified manufacturer’s protocol. In brief, live organoids were placed in an Ultra-Low Attachment 6-well plate, manually minced in a Papain+DNase solution in Earle’s medium, and incubated at 37 °C in an orbital shaker at 100 RPM for 30 minutes. Then, the pieces were gently broken up with a 1 mL pipette before returning to shaking in an incubator for an additional 20-30 minutes. Afterwards, using a 10 mL pipette, the solution with the minced-up pieces was gently mixed up and down 10 times and transferred to an empty 15 mL conical tube to allow undissociated debris to settle for ∼2 minutes. Subsequently, the cell suspension was transferred to a 3:8-diluted ovomucoid protease inhibitor solution in Earle’s medium, mixed, and then centrifuged at 300 x g for 7 minutes. The supernatant was then discarded, and the cells were gently resuspended in 1% BSA-supplemented basal medium before filtering through a 40 µm mesh filter. Cells were then counted and confirmed to have viabilities above 85% using a Countess II Cell Counter (Thermo-Fisher). Finally, samples were normalized to a concentration of ∼1000 cells/µL and transferred to 5 mL polypropylene tubes for processing at The Scripps Research Institute Genome Sequencing Core facility.

#### scRNA-seq

Using the Chromium Next GEM single cell 3’ Cellplex v3.1 and Chip G kits (10x Genomics), the single cell partitioning, cDNA preparation, and sequencing libraries were prepared according to the manufacturer’s protocol, using one 10x reaction for each experiment. After labeling the samples and amplifying the transcriptomic libraries, they were sequenced using on an Illumina Novaseq 6000 (SP or S1, depending on the run size). An average depth of ∼30,000 reads per cell was targeted. With a total number of 35,000 - 50,000 cells sequenced per run, adding up to 1.6 billion reads. See **Supplementary Table S1** for details.

#### scRNA-seq Data Preprocessing

The raw sequencer results were aligned to the GRCh38 genome assembly and de-multiplexed using Cell Ranger (10x Genomics) with default parameters. The aligned gene expression matrices were then processed on R via Seurat (ver.4.3.0.1).

First, potential dead cells and doublets were removed by removing any data points with mitochondrial gene load above 10% or with number of detected genes under 500 or above 7500. The data was then log normalized, after which 2500 highly variable genes were identified and data was scaled using the default LogNormalize, vst, and negbinom methods; respectively. PCA and Uniform Manifold Approximation and Projection (UMAP) embedding were then performed with default parameters using the top 35 principal components (PC). For a thorough removal of doublets, remaining doublets were identified using the DoubletFinder package. The proportion of doublet formation was estimated from the number of inter-sample doublets filtered by CellRanger for each independent experiment. After doublet removal, data was re-processed again with the same parameters. RNA counts and number of features across samples and unbiased clusters were assessed to ensure even sequencing quality and depth was maintained. Then, to correct for putative batch effects, dataset integration was performed^11^ (**Figure S2A**). Specifically, the data sets from each independent experiment were integrated based on their similar top 2000 highly variable features (HVFs) cluster profiles using the FindIntegrationAnchors and IntegrateData Seurat functions using default parameters. Dimensional reduction, clustering, and UMAP embedding were finally recalculated with the top 30 PC using the integrated data. The resulting integrated data set ensured neither experimental batch nor genotype contributed to more than 10% of any of the final unbiased clustering’s principal components.

#### scRNA-seq Data Annotation, and Validation

Following recommended single-cell annotation guidelines^12^, cell type annotation was done via the aggregate use of multiple unbiased annotation tools along with the gene enrichment within each unsupervised cluster (UC) to inform and validate their cell type identity based on their relative predicted cell type transcriptomic signatures. These tools included, SuperCT^13, 14^, suggesting single-cell-level annotation against reference cerebral organoid scRNAseq databases of Yoon et al.^5^ and Camp et al.^15^; and the scType tool^16^, providing unbiased cluster-level annotation tool based on an expanded set of canonical markers of all human developmental brain and central nervous system-related cell type markers assembled from the scType^16^, PanglaoDB^17^, and CellMarker^18, 19^ databases (See **Supplementary Table 2**). All clusters with agreement between these unbiased annotation tools were further validated by confirming they exhibited high and specific expression of a curated set of canonical markers-while also demonstrating minimal or absent expression of markers associated with other cell types-based on the aforementioned databases and the recently published CellxGene database (See **Supplementary Table 3**).^20^ Clusters with differing suggested identities were further sub-clustered and checked for their unbiased enriched genes (via Seurat’s FIndMarkers function) until clear agreement and validation was achieved for each subset. Any subcluster with still conflicting or low-confidence identities was annotated as unknown (“Unk”) under the “Other” general annotation. Finally, to facilitate data visualization, the negligibly small population of neuroepithelial cells (NEC) were merged into the transcriptionally similar choroid plexus-annotated “ChorPlx” label; similarly, given their small populations, oligodendrocyte progenitor cells (OPC) and oligodendrocyte (Olig)-annotated cells were merged into the “OPC/Olig” group. All automated and original annotations are available in the meta data. Visualization, cell type population, and gene expression plots were produced with standard Seurat package tools.

#### Gene Set Enrichment Analysis

We focused on the most relevant cell types in the COs; namely, excitatory neurons, inhibitory neurons, and astrocytes. Starting with the master fully annotated data set, each cell type was extracted at a time and Seurat’s FindMarkers function was used to calculate the differentially expressed genes (DEGs) between each genotype and their isogenic control with a minimum detection rate of 15% of the cells in either group and with a log fold change cut-off of 0.25.

#### Pseudotime Analysis

Trajectory inference and pseudotime analysis were conducted using Monocle3 (v1.3.7).^21^ Briefly, raw count matrices and metadata were extracted from the Seurat object to create a cell_data_set (CDS). Batch effect correction was applied using the align_cds() function. A principal graph representing the developmental trajectory was constructed using learn_graph() function. To ensure biological relevance and avoid artificial branching, clusters disconnected from the main developmental trajectory were manually inspected and excluded. The trajectory graph was recalculated following cluster removal. Pseudotime ordering was performed using order_cells() function, with root cells assigned based on early progenitor populations or specific clusters informed by validated annotation. The expression dynamics of key marker gene were visualized along pseudotime using plot_genes_in_pseudotime(), with additional facetting by cell types to reveal cell state-specific gene expression patterns. To investigate genotype-specific effects on trajectory, wild-type cells were subset based on genotype annotations and processed independently following the same analytical pipeline.

#### Microelectrode Array (MEA) Recordings of COs

COs of 6-8 weeks of age were attached to PEA/laminin–coated Axion Biosystems MEA plates and read on an Axion Biosystems Maestro Pro instrument. Specifically, 48-well Cytoview MEA plates (M768-tMEA-48B) were coated with a 0.05% Poly(ethyleneimine) (Sigma-Aldrich, P3143) in TC-grade water solution overnight at 37 °C. The wells were then thoroughly washed 3x with water, dried, and coated in a laminin I (R&D Systems, 3400-010-02) in PBS solution overnight at 37 °C. Just prior to use, the laminin solution was aspirated, and COs were transferred one per well to the plate. COs were left to attach at the center of each well for about 15 minutes at 37 °C before carefully filling the well with BrainPhys media (BrainPhys neuronal medium, 1x NeuroCult SM1, 1x N2 Max supplement). Two thirds of the media was changed every two days thereafter. The COs were allowed to adapt to this medium for at least 1 week before any data was recorded. To minimize metabolic and circadian rhythm activity variability, all CO electrophysiological activity data was collected for 5-minute epochs ∼23-24 hours after their last media change around the same time of day.

Analysis was performed using the Axion Biosystems software under the standard recommended settings for detection of action potentials on the Axion Navigator app and parameter analysis exported through the Neurometric Tool. Data from multiple timepoints was QC-cleaned to exclude electrode recordings without a minimum resistance value of 18 Ohms and/or without at least one voltage spike event per minute. Electrodes fulfilling these requisites were defined as “active electrodes”. A bursting event was defined by the default values of at least 5 spike fires within a 100 ms time interval. To prevent any bias caused by the number of active electrodes producing quality data, only weighted parameters were used, utilizing only or normalizing to the number of active electrodes. All timepoint recordings were aggregated using simple custom-made R scripts (available upon request) to calculate each CO’s average bursting rate and imported into GraphPad Prism for plotting.

#### CO Treatments

To carefully control conditions, individual COs were placed into separate Ultra-Low Attachment 24-well plate wells with a pre-defined media volume (e.g. 1.5 mL) across all conditions. DMSO vehicle was normalized to be equal across all conditions independent of their compound concentration. COs were treated on every media change feed every two days in, shaking in orbital shakers at 100 RPM for experiments meant for immunoblotting or immunofluorescence analysis. Cellular viability was complementarily assessed with high-sensitivity lactate dehydrogenase (LDH)-Glo kits following manufacturers’ recommendations. For large-scale experiments, samples were diluted 1:25 in LDH Storage buffer (200 mM Tris-HCl, 10% Glycerol, 1% BSA, 7.3 pH) and kept at -80C until ready to assay all timepoints together. For electrophysiology experiments in MEA, basal measurements were taken for at least 1 week prior to treatment. Then, at the time of treatment, the compounds were diluted in fresh BrainPhys media at concentrations accounting for the dilution due to partial media change. MEA viability was followed in real time throughout the whole experiment through the viability module (via the electrode resistances) collected at each activity recording. During the treatment period, more frequent recordings were performed during the previously established stable activity period of 12-to-28 hours post-media change. The Recovery period was defined as that after the compound wash out through 5x consecutive fresh BrainPhys media replacements. To assess drug effects, mean activity values were calculated for each individual CO from the multiple recordings during 1-1.5 weeks prior to treatment (“Basal”), 12-28 hours under drug exposure (Treatment), and 24 to 72 hours after drug wash out (still changing media to record only during the aforementioned stable activity periods).

#### Image Analysis

Images were processed using ImageJ (Fiji v.2.9.0/1.53t). For CO size measurements, Cell Magic Wand tool was used to select each CO contour before measuring area. Regarding surface area coverage calculations, after which the area function measurement was used to quantify the percent surface area from individual images. Regarding puncta count, a mask identifying particles by size was used to filter and count distinct puncta from images. Measured signal from each image used for analysis was normalized to the total number of nuclei. Individual cells were identified by segmenting Hoechst-stained images on the DAPI channel using a nuclei mask. This mask filtered nuclei by size and separated conjoined nuclei with the watershed function. Any artifacts or images with unreliable morphological staining, arising from technical issues or tissue anomalies such as folds or holes, were omitted from analyses.

### QUANTIFICATION AND STATISTICAL ANALYSIS

Data is reported as means ± standard deviation. Each dot in the figures represents biological replicate samples of individual CO(s), unless otherwise stated in the figure legend. Experimental replicates were defined as biological replicates with samples deriving from both, different CO induction date and the date when the assay was performed. Number of samples per group (n) represents biological replicates and is indicated in legends. In Aβ ELISA, LDH-Glo, and MEA recordings duplicate to triplicate measurements of individual COs, considered technical replicates, were averaged to use their mean value as single biological sample data points. All data comparisons analyses were done in GraphPad Prism for Mac (version 9 through 10.0.3) using analysis of variance (ANOVA) analysis with the Prism-recommended post-hoc multiple comparisons test correction, as indicated in figure legends. Welch’s ANOVA was used where indicated in figure legends when data was statistically shown to be heteroscedastic. Western blot (WB) data with experimental replicates was analyzed by 2-way ANOVA with genotype/condition and experimental batch as the fixed factors and with Dunnett’s multiple comparisons correction post-hoc test. When possible, WB data was normalized by inter-blot controls, as indicated in figure legends.^23^ In cases where data was found not to be normally distributed, data was log-transformed and re-tested for normality; any such data is plotted in its log-transformed form in the figures and indicated in the legends. Data with any overt outliers were tested via Grubbs’s test with a conservative alpha of 0.01. If confirmed by test, only up to a single outlier per data set was removed. When analyzing changes in CO functional activity due to treatments, pair-wise comparisons across timepoints were assessed using a linear mixed-effect model with restricted maximum likelihood (REML) estimation to account for repeated measures and within-organoid correlations. Subjects (organoid replicates) were treated as random effects, enabling unbiased variance estimation and robust inference for changes between matched timepoints without sphericity assumptions. P-value significance was illustrated according to: *p<0.05, **p<0.01, ***p<0.001, ****p<0.0001.

## ACKNOWLEDGMENTS

The hiPSC isogenic set lines 1 and 2 were provided by Larry Goldstein (UC San Diego) and Marc Tessier-Lavigne (Rockefeller University and Stanford University), respectively. We want to thank Jessica E. Young for kindly providing us with the WT/WT (2) control cell line. We want to acknowledge Henry Scott, Nhi Buu Lang for assisting in the maintenance of some organoid cultures; Jorryn Wu and Mingi Kim for experimental insights helpful to the project; Hongfan Peng, Juan Piña-Crespo, Xu Zhang, and Scott R. McKercher, for facilitating resources critical to the project; the Scripps Research Center for Computational Biology and Bioinformatics (CCBB) core, for facilitating raw data output and resources for scRNA-seq data processing; Alan Chu, for his support in using Calibr’s MSD instrument for the beta amyloid measurements; and Emily P. Bentley, for proofreading, editing, and improving the manuscript writing. We are grateful to Sneha Arora, Evan Powers, Luke Wiseman, and John Yates III at Scripps Research; and Kristin Baldwin at Columbia University for their helpful feedback during the project. This study was made possible by support in part by NIH grants (R01 AG073418 to J.W.K., U01 AG088679, R01 AG056259, R35 AG071734 to S.A.L., and TL1 TR002551 to S.R.L.), the Freedom Together Foundation to J.W.K., and the ARCS Foundation to S.R.L.

## AUTHOR CONTRIBUTIONS

Project conceptualization, supervision, and administration, S.R.L., J.W.K., and S.A.L.; cerebral organoid protocol optimization, S.R.L. and N.D.; cell and organoid culture, S.R.L., N.D., J.P.S, C.C.K., C.B, J.C., M.A., and A.B.; biochemistry, S.R.L., C.C.K., M.A., and A.P.; immunohistochemistry, S.R.L., A.P., J.P.S., C.C.K., and M.B.; microscopy, A.P, S.R.L., J.P.S., and C.C.K.; organoid electrophysiology and analysis, S.R.L., S.G., M.T., S.A.L.; single-cell RNA-seq design and preparation, S.R.L., D.T., T.S.M., S.R.H.; scRNA-seq analysis, S.R.L., Y.W., W.L., N.J.S.; mass spectrometry, Z.G.; writing of original draft, S.R.L.; manuscript proof-reading and revisions, S.R.L., A.B., M.B., J.W.K, and S.A.L.

