## Supplemental Figures S1-S5 for "Autophagy activators normalize aberrant Tau proteostasis and rescue synapses in human familial Alzheimer’s disease iPSC-derived cortical organoids"

Fig.S6

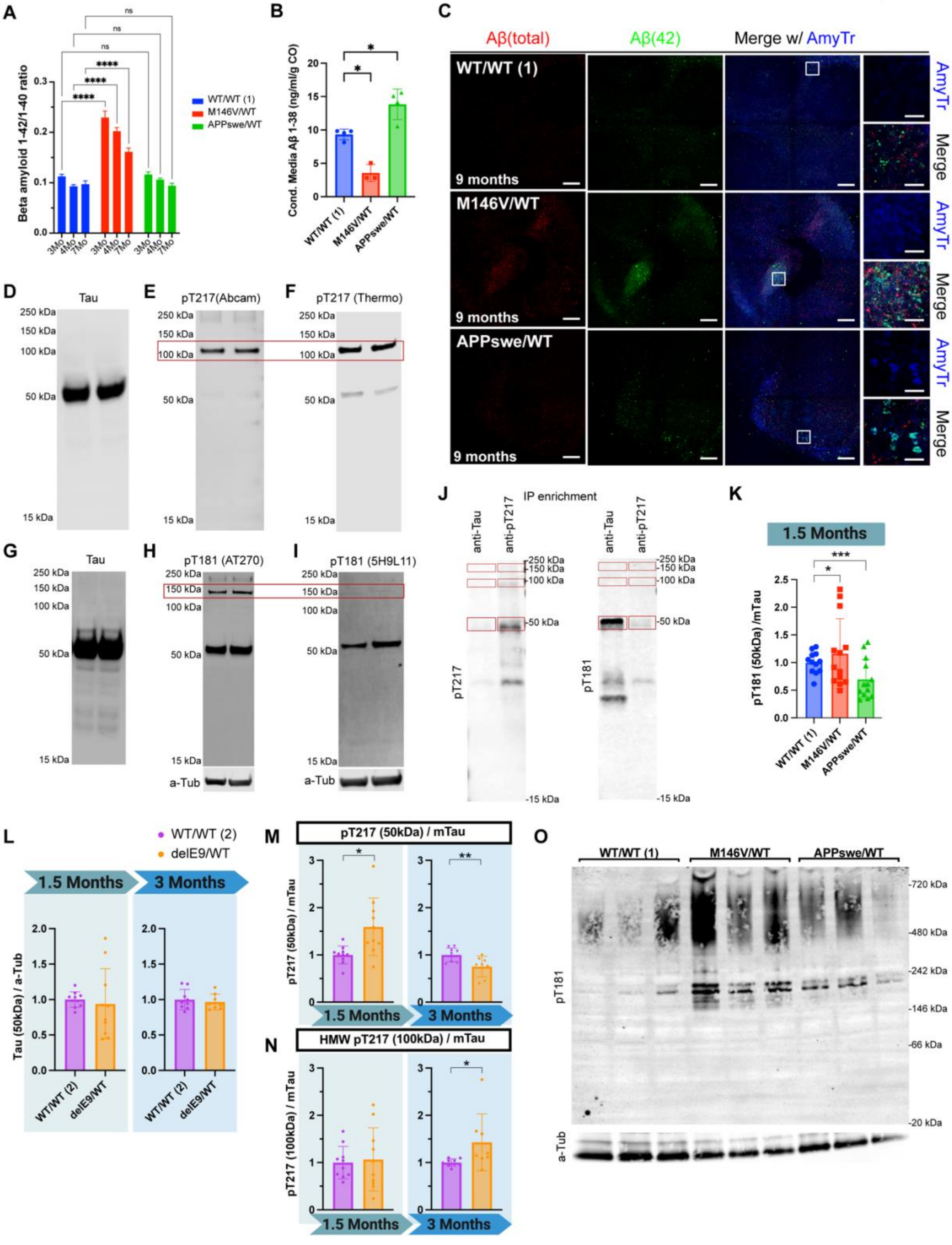
A
Beta amyloid 1-42/1-40 ratio
WT/WT (1)
M146V/WT
APPswe/WT
ns
****
3Mo
4Mo
7Mo
B
Cond. Media Aβ 1-38 (ng/ml/g CO)
WT/WT (1)
M146V/WT
APPswe/WT
*
C
Aβ(total)
Aβ(42)
Merge w/ AmyTr
WT/WT (1)
M146V/WT
APPswe/WT
9 months
AmyTr
Merge
D
Tau
E
pT217(Abcam)
F
pT217 (Thermo)
250 kDa
150 kDa
100 kDa
50 kDa
15 kDa
G
Tau
H
pT181 (AT270)
I
pT181 (5H9L11)
a-Tub
J
IP enrichment
anti-Tau
anti-pT217
pT217
pT181
K
1.5 Months
pT181 (50kDa) /mTau
L
WT/WT (2)
delE9/WT
1.5 Months
3 Months
Tau (50kDa) / a-Tub
M
pT217 (50kDa) / mTau
N
HMW pT217 (100kDa) / mTau
pT217 (100kDa) / mTau
O
WT/WT (1)
M146V/WT
APPswe/WT
720 kDa
480 kDa
242 kDa
146 kDa
66 kDa
20 kDa
pT181
a-Tub

### Supplementary Fig. S6: Additional validation of Aβ and pTau pathology assessments in the COs, related to Figure 3.

(A) Ratio in CO-conditioned media of $A\beta_{1-42}$ to $A\beta_{1-40}$ at 3, 4, and 7-month timepoints. n = 3-4 COs per condition.
(B) $A\beta_{1-38}$ peptide concentration in conditioned media from 5-6 months-of-age COs for isogenic set 1. Concentrations were below quantifiable levels for COs from isog. set 2.
(C) Immunostaining of the isogenic COs at 9-month timepoint with the amyloid stain, AmyTracker (AmyTr), and a total Aβ antibody. Scale bar, 100 µm.
(D-F) WB of two $PSEN1^{delE9/WT}$ 1.5 months-of-age CO lysates probed with the pan-Tau antibody (mTau) (D) or with p217 antibody (Thermo AT270, RRID: AB_223651) after incubation with a saturating concentration of pan-Tau antibody (E). (F) Replicate WB of the same CO lysates probed with a different phospho-Tau T217 antibody, clone 5H9L11 (Thermo, RRID: AB_2532491). Red box highlights the consistent HMW band persisting in the reduced SDS-PAGE.
(G-I) WB of two independent lysates from $PSEN1^{delE9/WT}$ 1.5-month timepoint COs probed with the pan-Tau antibody (mTau) (G) or with pT181 antibody (Thermo AT270, RRID: AB_223651) after incubation with a saturating concentration of pan-Tau antibody (H). (I) WB replicate of the same CO lysates probed with a different phospho-Tau T181 antibody, clone 5H9L11 (Thermo, RRID: AB_2532491). Red box highlights the consistent HMW band persisting in the reduced SDS-PAGE.
(J) WB of immunoprecipitate (IP) of 3 months-of-age $PSEN1^{M146V/WT}$ CO lysate to enrich for either Tau or pT217. Red boxes indicate the bands directly assessed by mass spectrometry. Tau (MAPT) was identified in 5 out of 6 samples (all except one of the 105kDa samples)
(K) WB Quantification of phospho-Tau pT181 in CO lysates at the 1.5-month timepoint, as normalized to mTau. n = 10-14 COs per genotype from 2-4 independent experiments.
(L) WB Quantification of monomeric Tau (mTau) in isogenic set 2 CO lysates at the 1.5- (*left*) and 3-month (*right*) timepoints, as normalized to α-Tubulin (a-Tub). n = 8-10 COs per genotype from 3 independent experiments per timepoint.
(M-N) WB Quantification of phospho-Tau pT217 at its monomeric (~50 kDa)(M), and strongest (~100 kDa)(N) bands in isog. set 2 CO lysates at the 1.5 and 3-month timepoints, as normalized to mTau. n = 8-10 COs per genotype from 3 independent experiments per timepoint.
(O) Representative native PAGE of 4-month timepoint CO lysates probed for pT181 and α-Tubulin.
Data are mean ± SD. Analyses by ANOVA with Dunnett's post-hoc test.

SUPP. FIG. S7

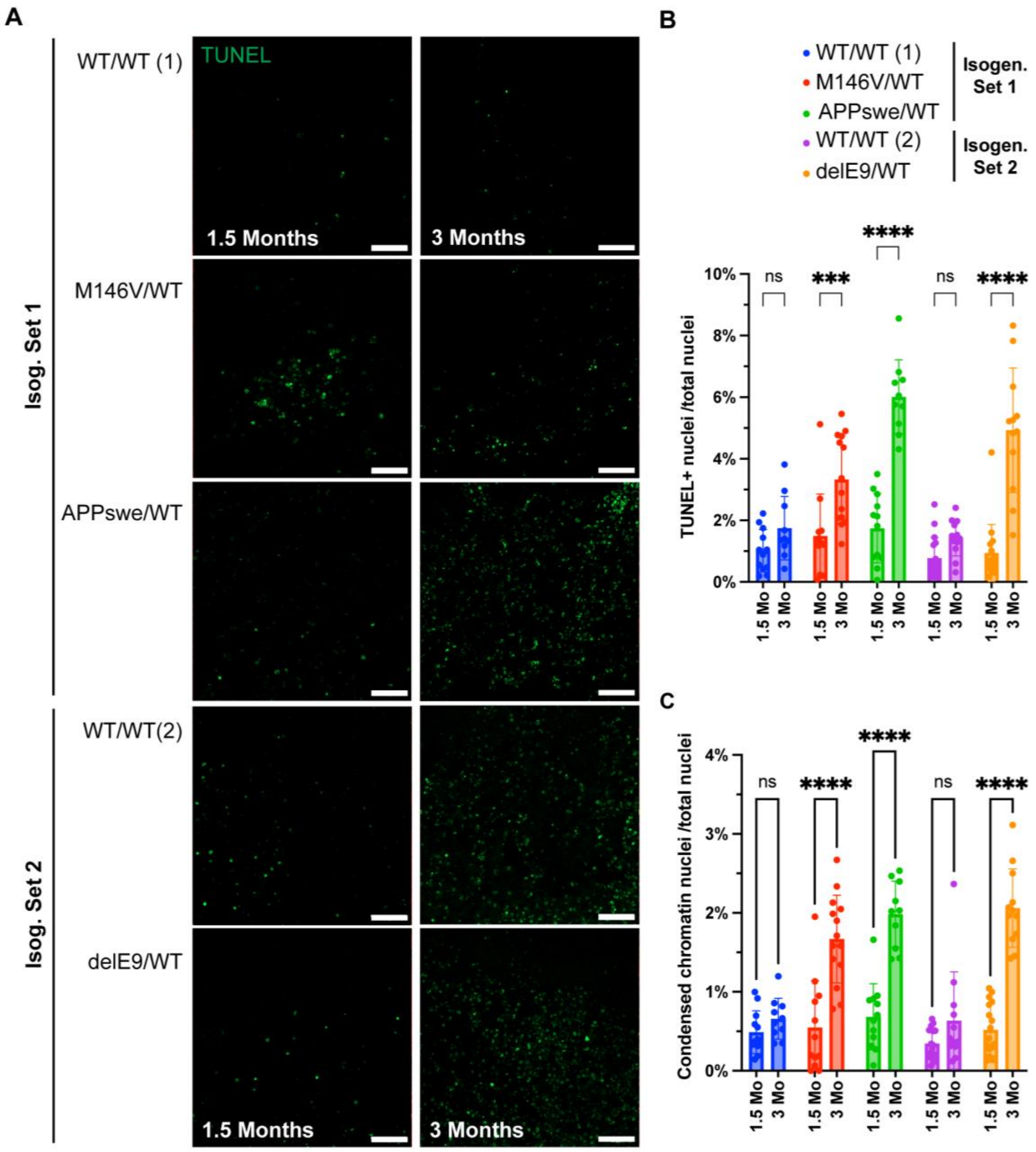

Supplementary Fig. S7: AD COs display signs of neurodegeneration, related to Figure 3.

(A) Representative images of 6 weeks and 3 months-of-age COs for TUNEL (Terminal deoxynucleotidyl transferase dUTP nick end labeling) staining for detecting apoptotic and/or necroptotic cells. Scale bar, 50 µm.

(B and C) Independent quantification of either the total number of TUNEL+ nuclei (B) or the number of nuclei with condensed chromatin (C), normalized to the total number of nuclei. Total number and chromatin condensation counts were both assessed by overexposed post-fixation staining with Propidium Iodide to visualize all nuclei, n = 12-17 images from at least 3 COs per condition. Data are mean ± SD. Analysis by ANOVA with Sidak's post-hoc test.

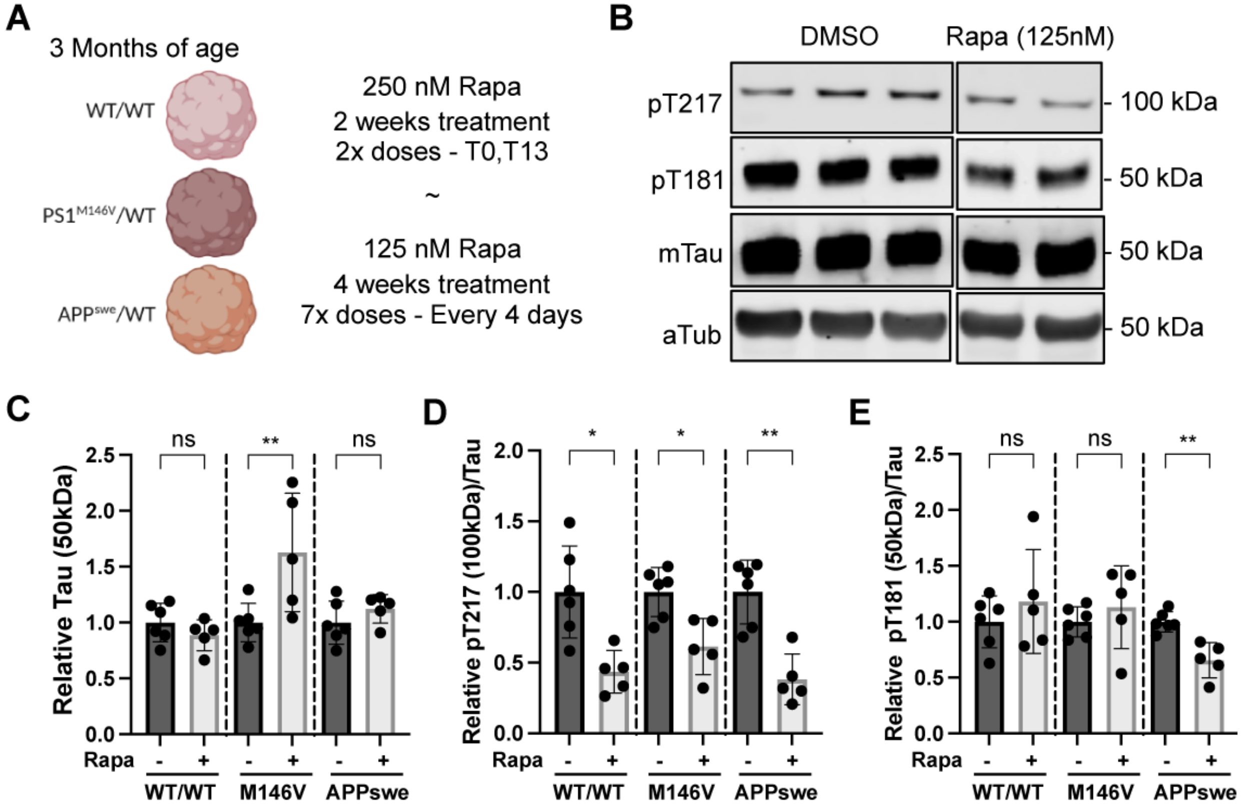

Supplementary Fig. S8: Chronic mTOR-dependent autophagy activation reverses AD-associated pTau pathologies, related to Figure 4.

(A) Diagram describing the experimental set-up of the 2-week-long and 4-week-long mTOR-mediated Rapamycin (Rapa) treatment of 3-month timepoint (by end of treatment) COs.

(B) Representative WB of CO lysates at the end of the 2-week-long treatment probed for their Tau and relative pTau abundances.

(C-E) WB Quantification for the normalized monomeric Tau (C), pT217 (~100kDa) (D), and pT181 (~50 kDa) (E) abundances after the treatment. Bands at other molecular weights were too low to quantify. n = 5-6 COs per genotype per condition from 2 independent experiments. Data are mean ± SD. Analyses by ANOVA with Dunnett’s post-hoc test.

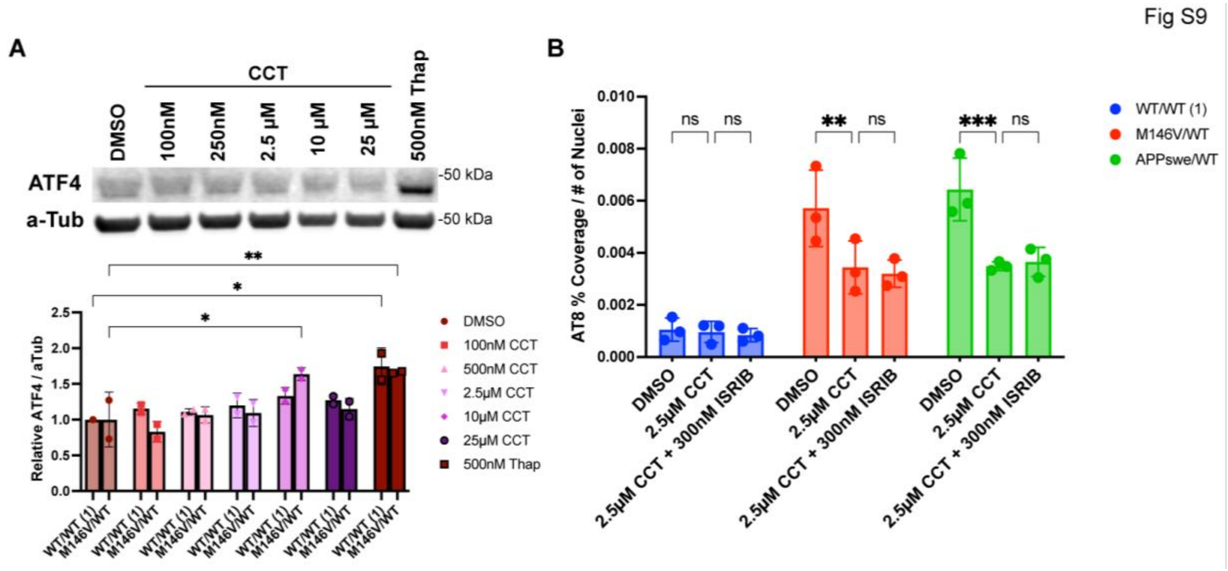

### Supplementary Fig. S9: CCT treatment does not activate or rely on PERK at concentrations used in the study, related to Figure 5.

(A) Representative WB of lysed COs after 6-day treatment with DMSO vehicle, 100 nM – 25 µM CCT, or 500 nM Thapsigargin (as a PERK activator positive control) probed for ATF4. COs were treated on day (D)0, and D5, and lysed 24 hours after the second treatment day. Quantification of ATF4 signal normalized to α-tubulin (a-Tub) and relative to DMSO value.
(B) Quantification of the paired helical filament (PHF) aggregated pTau ($AT8^+$) percent coverage normalized to nuclei number after 2-week treatment with 2.5 µM CCT with or without 300 nM ISRIB (integrated stress response inhibitor). n = 3 COs per condition. Data are mean ± SD. Analyses by ANOVA with Dunnett's post-hoc test.

Fig S10

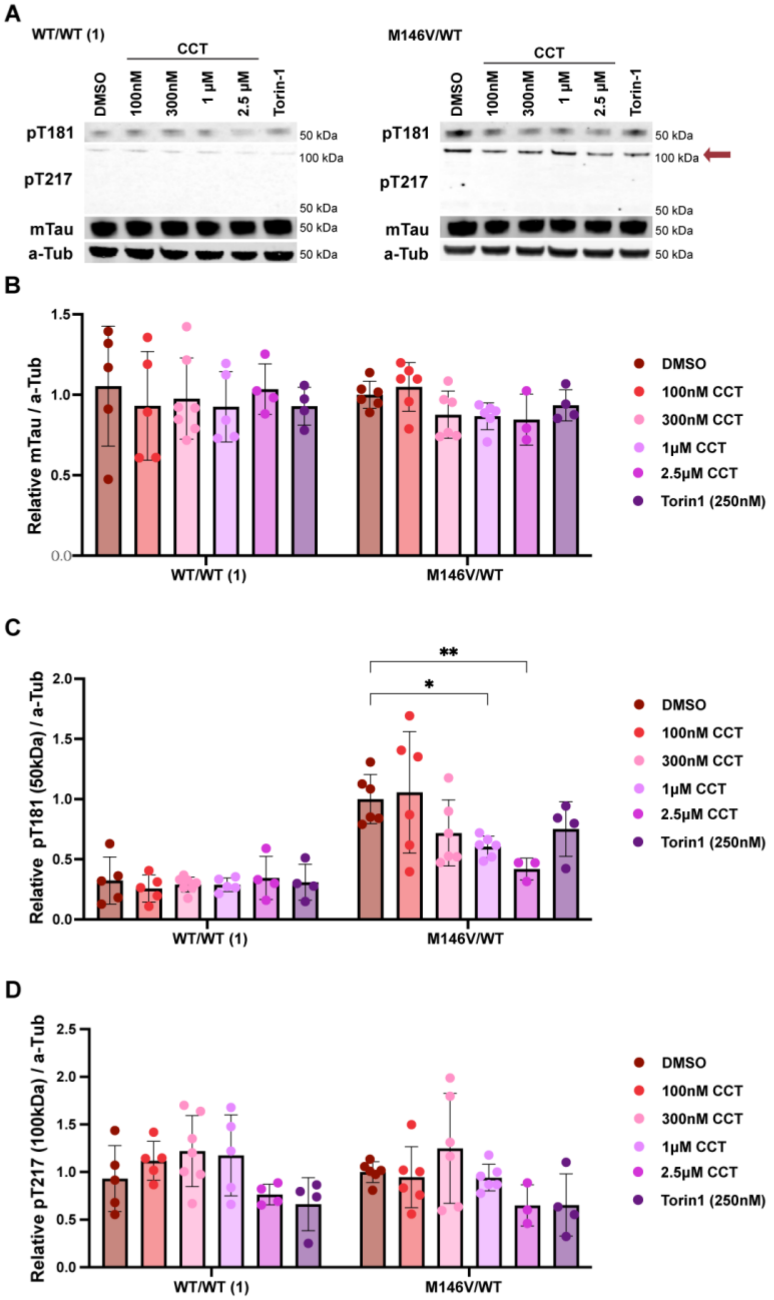

### Supplementary Fig. S10: Chronic autophagy activation differentially reduces AD-associated pT217 and pT181, related to Figure 5.

(A) Example WBs of the WT/WT (1) and PSEN1 M146V/WT COs probed for monomeric Tau (mTau), phosphorylated Tau 181 (pT181), phosphorylated Tau 217 (pT217), and α-tubulin (a-Tub) at the 3-month timepoint after a 4-week treatment with DMSO vehicle, CCT, or Torin-1. (B-D) WB Quantification of the WT/WT (1) and PSEN1 (PS1(M146V)) mutant CO lysates for a-Tub-normalized mTau (B), pT181 (C), and pT217 (D) at the end of a 6-week-long treatment. Data from multiple experiments are normalized to mean PS1(M146V) DMSO values via inter-blot controls. n = 12-23 COs per genotype from 2 independent experiments. Data are mean ± SD. Analysis by ANOVA with Dunnett's post-hoc test.

FIG.S11

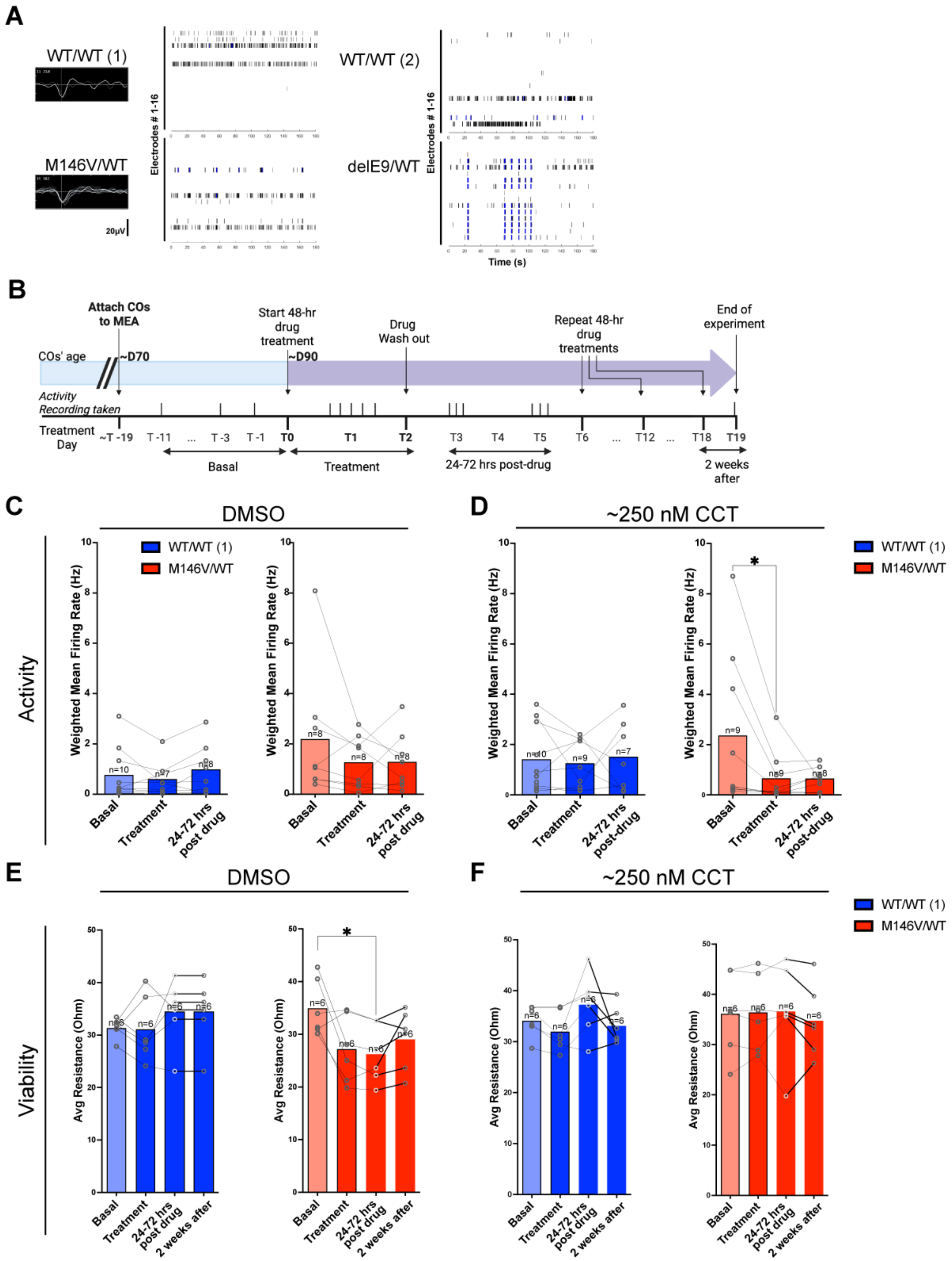

A
WT/WT (1)
M146V/WT
20μV
Electrodes # 1-16
WT/WT (2)
delE9/WT
Time (s)
B
Attach COs to MEA
Start 48-hr drug treatment
Drug Wash out
Repeat 48-hr drug treatments
End of experiment
COs' age
~D70
~D90
Activity Recording taken
Treatment Day
~T -19
T -11
T -3
T -1
T0
T1
T2
T3
T4
T5
T6
T12
T18
T19
Basal
Treatment
24-72 hrs post-drug
2 weeks after
C
DMSO
D
~250 nM CCT
Activity
WT/WT (1)
M146V/WT
Weighted Mean Firing Rate (Hz)
Basal
Treatment
24-72 hrs post drug
E
DMSO
F
~250 nM CCT
Viability
Avg Resistance (Ohm)
2 weeks after

### Supplementary Fig. S11: AD COs display neuronal hyperexcitability that is dose-dependently reduced by CCT, related to Figure 5.

(A) Representative raster plots and mean firing rate (Hz) quantification over the span of 180 seconds of ~3 months-of-age of PSEN1 fAD COs from isogenic sets 1 and 2.
(B) Schematic describing treatment with CCT and activity recording of TL (isogenic set 1) COs.
(C-D) Quantification of WT/WT and PSEN1$^{M146V}$ CO neuronal activity, as calculated by the weighted mean firing rate (Hz) during basal, treatment, and recovery periods (the latter obtained 1-3 days after washout) for DMSO (C) and ~250 nM CCT (D). Number of COs recorded indicated above bars in the figure.
(E-F) Quantification of WT/WT and PSEN1$^{M146V}$ COs' viability, as reflected by mean electrode resistance (in kOhms) during basal, treatment (data obtained 12-24 hours after first dose), and recovery periods (data obtained 1-3 days after washout) for the DMSO (F) and ~250 nM CCT (G) conditions. Data are mean ± SD. Number of individual COs recorded (from 2 independent experiments) indicated above bars in the figure. Analysis by Mixed-effect model (RELM) for matched repeated measures.
