## Supplemental Figures S6-S11 for "Autophagy activators normalize aberrant Tau proteostasis and rescue synapses in human familial Alzheimer’s disease iPSC-derived cortical organoids"

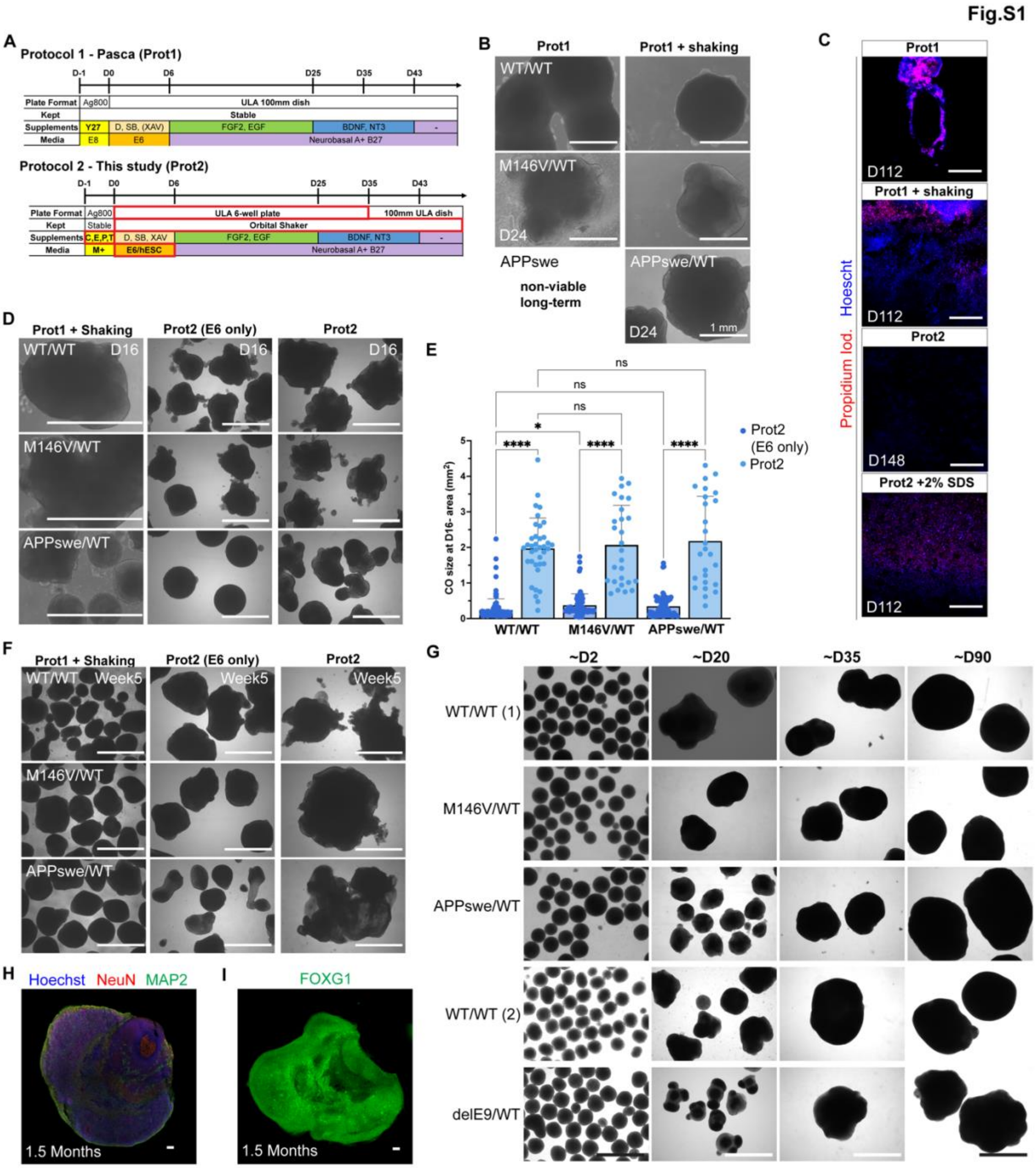

### Supplementary Fig. S1: Protocol modifications to generate reproducible, healthy COs across all genotypes, related to Figure

(A) Schematic summarizing the key modifications (Protocol 2, highlighted in red) made over the method of Pasca (Protocol 1). M+: mTeSR plus; C,E,P,T: Chroman 1, Emricasan, Polyamines, Trans-ISRIB; E6/hESC: 50/50 mix of Essential 6 and hESC media. See STAR Methods for details.

(B) Representative bright field images of typical organoids maintained in 10 cm dishes (*left*) vs. in 6-well plates in an orbital shaker at day (D)24 (*right*). Scale bar, 1 mm.

(C) $APP^{swe}$ COs live-stained with propidium iodide and imaged cross-sectionally after cutting in half to show their relative degree of necrosis across different protocol iterations. As positive control, one of the COs was pre-conditioned in media containing 2% SDS. Scale bar, 500 µm.
(D) Representative bright field images showing the size and heterogeneity of isogenic sets of COs at D16 depending on the induction method. E6: Essential 6 medium (Gibco); Prot2: 50/50 mix of E6 and hESC medium.
(E) Quantification of the mean CO size (maximal cross-sectional area) at D16 comparing the effect of the induction media, n = 77-109 COs per genotype for "E6 only", n = 26-37 COs per genotype for E6/hESC; each condition from 4 independent CO induction experiments. Data are mean ± SD. Analysis by Welch's ANOVA with Dunnett's T3 post-hoc test.
(F) Representative bright field images showing the size and heterogeneity of isogenic sets of COs at approximately D35 (Week 5) depending on the induction medium used.
(G) Representative brightfield images of COs of each cell line produced with Protocol 2, at ~2, 20, 39, and 90 days in culture. Quantification shown in Figure 1B. Scale bar, 2 mm.
(H) Representative immunofluorescent (IF) staining of mature postmitotic neurons ($MAP2^+$, $NeuN^+$) in COs at ~6-week timepoint. Scale bar, 100 µm.
(I) Representative IF stain for forebrain ($FOXG1^+$) fate in COs at 1.5-month timepoint. Scale bar, 100 µm.

Fig.S2

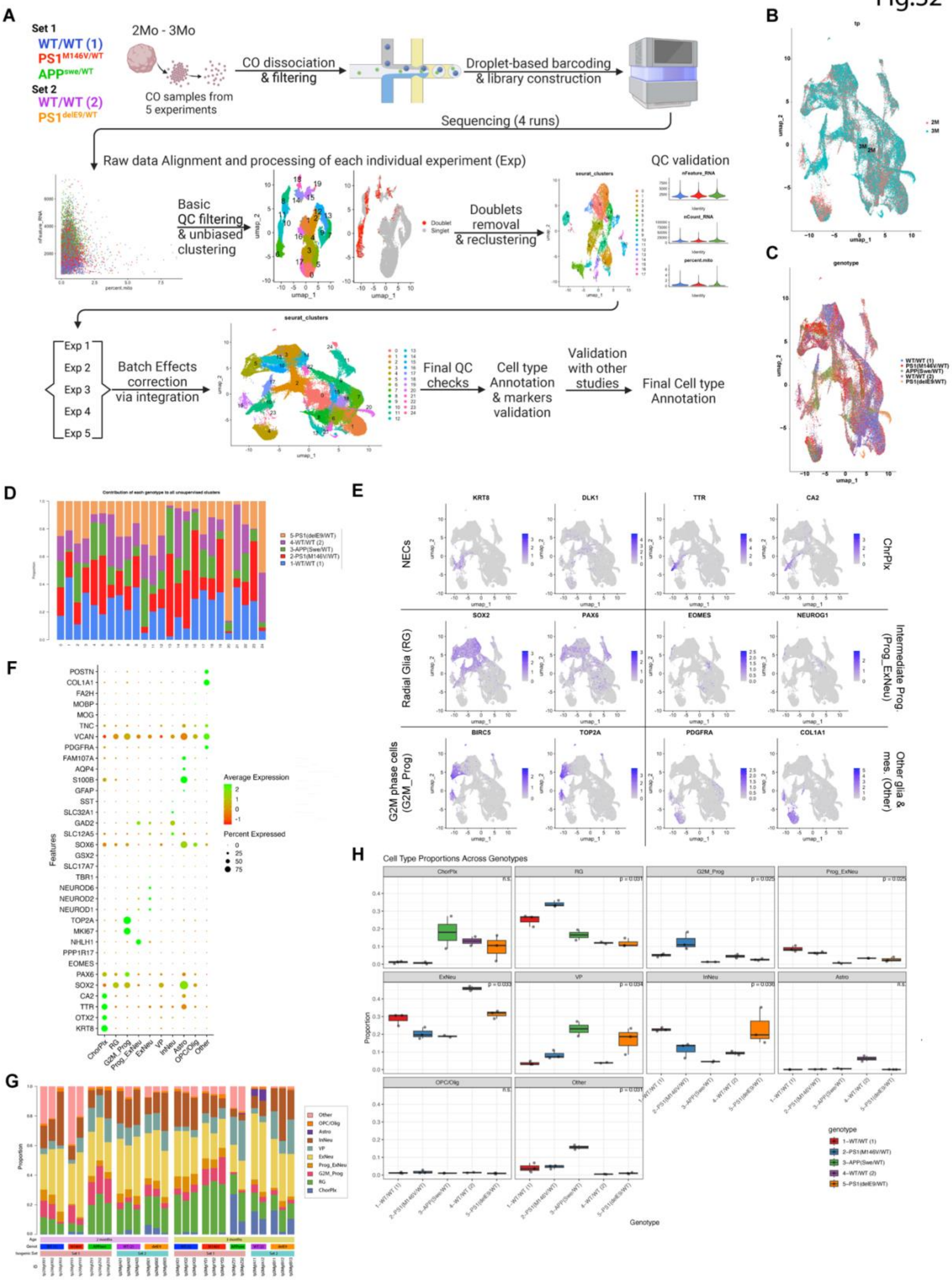

A
Set 1
WT/WT (1)
PS1M146V/WT
APPswe/WT
Set 2
WT/WT (2)
PS1delE9/WT
2Mo - 3Mo
CO samples from 5 experiments
CO dissociation & filtering
Droplet-based barcoding & library construction
Sequencing (4 runs)
Raw data Alignment and processing of each individual experiment (Exp)
QC validation
Basic QC filtering & unbiased clustering
Doublet
Singlet
Doublets removal & reclustering
Exp 1
Exp 2
Exp 3
Exp 4
Exp 5
Batch Effects correction via integration
Final QC checks
Cell type Annotation & markers validation
Validation with other studies
Final Cell type Annotation
B
C
D
E
NECs
ChrPlx
Radial Glia (RG)
Intermediate Prog. (Prog_ExNeu)
G2M phase cells (G2M_Prog)
Other glia & mes. (Other)
F
Average Expression
Percent Expressed
Features
G
H
Cell Type Proportions Across Genotypes
Genotype

### Supplementary Fig. S2: scRNA-seq cluster annotation and validation, related to Figure 2.

(A) Diagram describing the sample and data processing of 5 experiments for the COs' scRNA-seq along the respective UMAP clustering plots of the data before and after integration. The final set contained 70,854 high quality cells. See STAR Methods for details.
(B) UMAP cluster indicating the cell type mutant genotypes distribution (showing how they are similarly contributing to most clusters) along a breakdown of each gene mutant by their timepoint. 2-month in red; 3-month in aqua.
(C) UMAP cluster indicating the cell type mutant genotypes distribution (showing how they are contributing to all UMAP cell clusters.
(D) Bar plot breakdown showing an alternative representation of all CO genotypes contributing to all of the Seurat unbiased clusters.
(E) Feature plots displaying the localized expression of hallmark markers for other identified CO cell types not already displayed in Fig.2B. Highlighted here are KRT8 and DLK1 for neuroepithelial cells (NECs); TTR and CA2 for choroid plexus (ChrPlx); SOX2 and PAX6 for Radial glia (RG); EOMES and NEUROG1 for intermediate progenitors (Prog_ExNeu); BIRC5 and TOP2A for cells in G2-M phase undergoing mitosis (G2M_Prog); and PDGFRA and COL1A1 for neural crest Schwann-like glia and mesenchymal progenitors (Other).
(F) Dot plot of cell type and cell state hallmark genes across the annotated scRNA-seq groups to supplement Fig. 2C.
(G) Proportions of all major cell types for each individual CO sample sequenced, organized by differentiation age stage, genotype, and experimental batch.
(H) Box and whisker plots of the proportional representation of the different cell types in the 3-month timepoint COs across all genotypes. Analysis for each cell type by Kruskal-Wallis Rank Sum test.

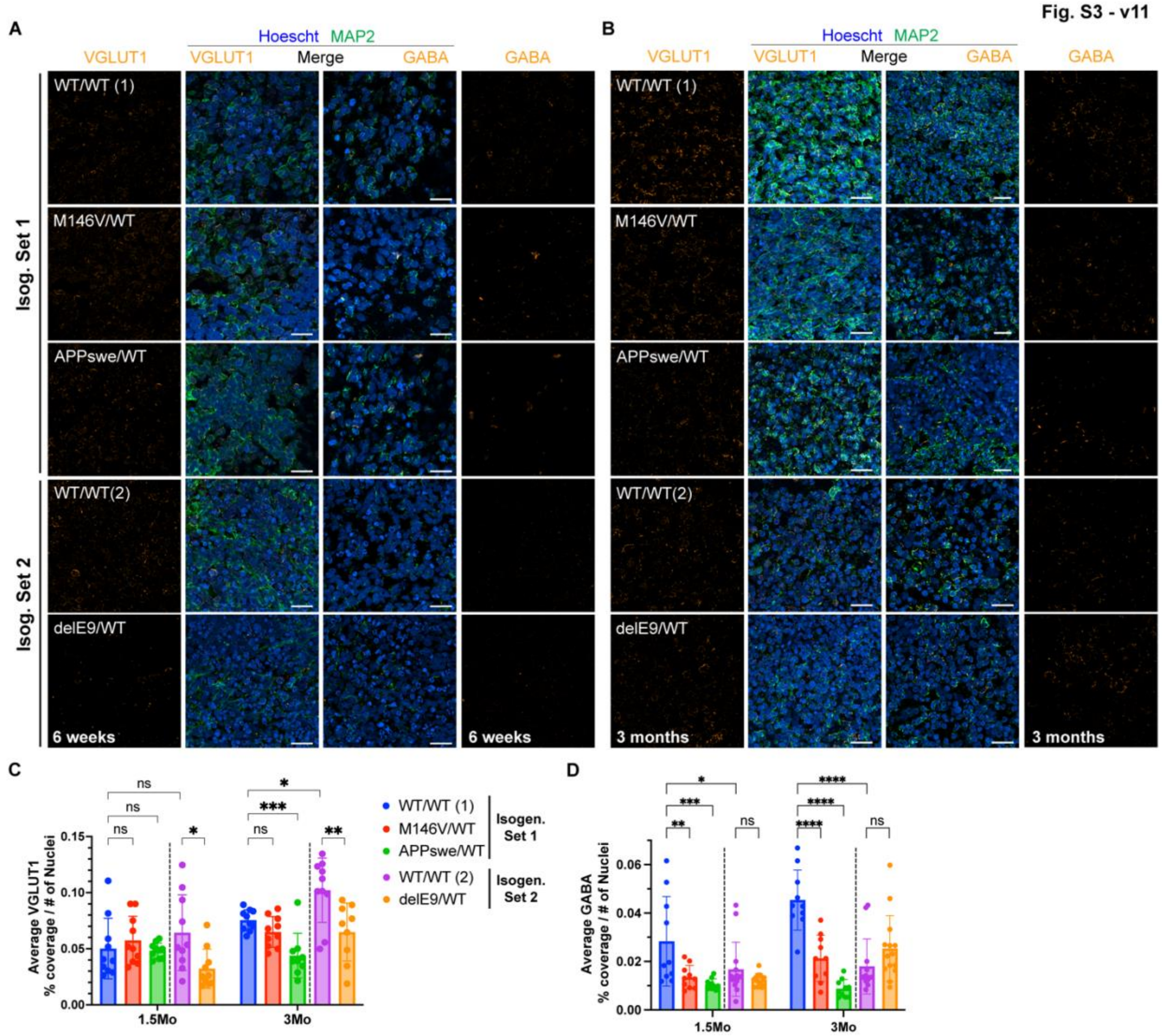

Supplementary Fig. S3: CO neuronal immunofluorescent validation, related to Figure 2.

(A and B) Representative Immunofluorescence (IF) images of 6-week (A) and 3-month (B) timepoint COs for VGLUT1 (Excitatory neurons, *left*) and GABA (Inhibitory neurons, *right*) with Hoechst and MAP2 as pan-nuclear and mature postmitotic neuronal counterstains. Scale bar, 20 µm.

(C and D), Quantification of VGLUT1+ (C) and GABA+ (D) relative proportion as percent coverage area normalized to number of nuclei for 1.5- and 3-month timepoint COs, n = 9-10 images from 3-5 COs per condition per timepoint. Data are mean ± SD. Analysis by ANOVA with Dunnett's post-hoc test.

Fig.S4

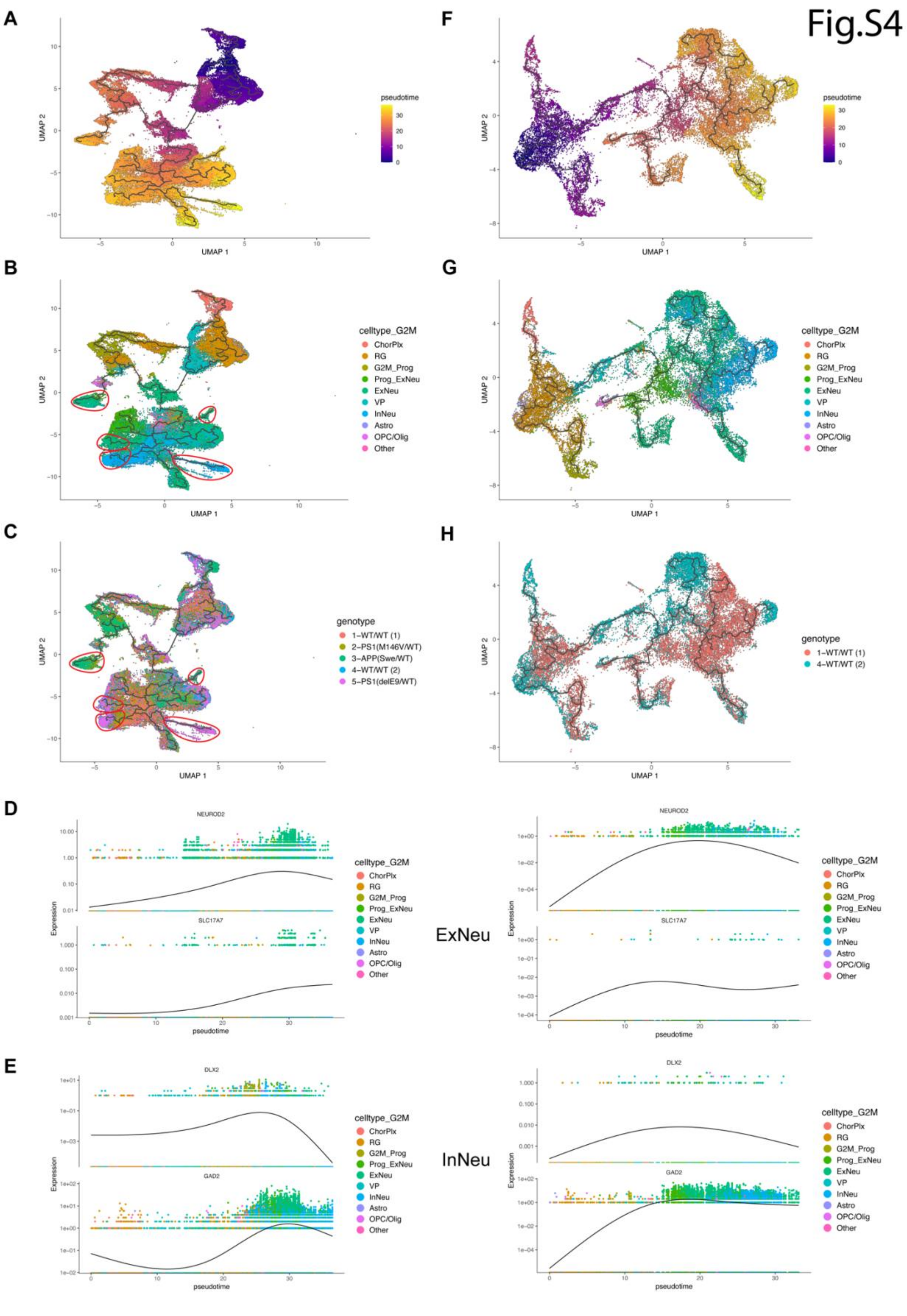

A
B
C
D
E
F
G
H
UMAP 1
UMAP 2
pseudotime
celltype_G2M
ChorPlx
RG
G2M_Prog
Prog_ExNeu
ExNeu
VP
InNeu
Astro
OPC/Olig
Other
genotype
1-WT/WT (1)
2-PS1(M146V/WT)
3-APP(Swe/WT)
4-WT/WT (2)
5-PS1(delE9/WT)
NEUROD2
SLC17A7
DLX2
GAD2
Expression
ExNeu
InNeu

### Supplementary Fig. S4: Pseudotime analysis reveals mutant-specific maturation trajectories of neural lineages, related to Figure 2.

(A-C) UMAP visualization showing the inferred developmental progression of all 5 genotypes along CO lineage differentiation. Colored by pseudotime (A); colored by cell type annotation (B); colored by genotype (C).
(D) Pseudotime plots of gene expression dynamics for immature ExNeu (NEUROD2) and mature ExNeu (SLC17A7), marking ExNeu lineage progression. All genotypes (*left*); WT/WT (from isogenic set 1) and WT/WT (from isogenic set 2) (*right*).
(E) Pseudotime plots of gene expression dynamics for immature InNeu (DLX2) and mature InNeu (GAD2), marking InNeu lineage progression. All genotypes (*left*); only WT/WT (1) & WT/WT (2) genotypes (*right*).
(F-H) UMAP visualization showing the inferred developmental progression of cells from the WT/WT genotypes (1 & 2) along CO lineage differentiation. F: colored by pseudotime; G: colored by cell type annotation; H: colored by genotype.
RG: radial glia; Prog_ExNeu: excitatory neurons progenitors; ExNeu: excitatory neurons; InNeu: inhibitory neurons; Astro: astrocytes; OPC/Olig: oligodendrocyte progenitors/oligodendrocytes; ChorPlx: choroid plexus cells.

Fig.S5

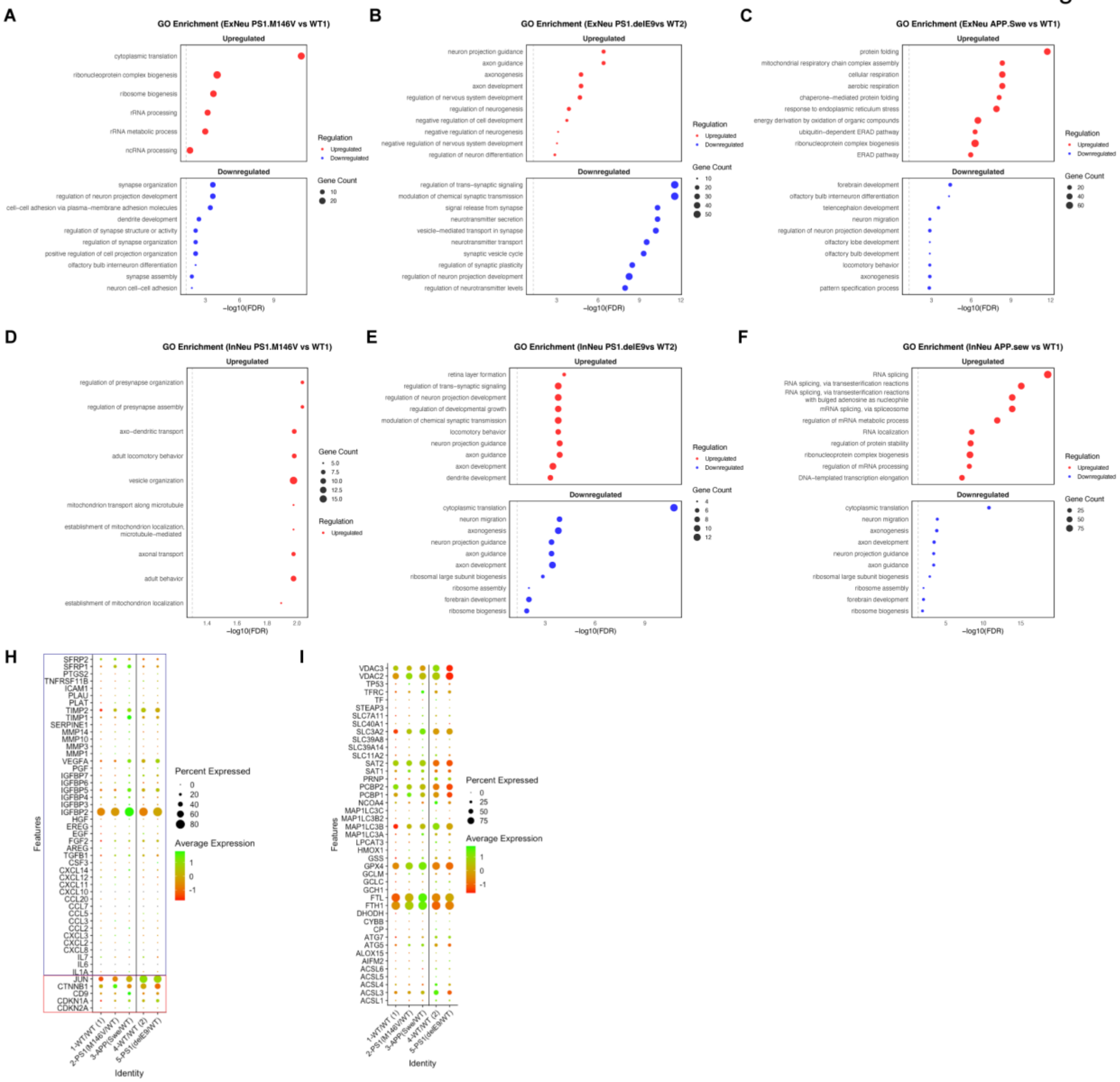

Supplementary Fig. S5: Transcriptomic analysis of AD vs WT COs, related to Figure 2.

(A-C) GO Enrichment in Excitatory neurons (ExNeu) for M146V/WT (A) delE9/WT (B) and APPswe/WT (C) vs their respective isogenic WT genotypes.

(D-F) GO Enrichment in Inhibitory neurons (InNeu) for M146V/WT (D) delE9/WT (E) and APPswe/WT (F) vs their respective isogenic WT genotypes.

(H) Dot plot of Senescence markers pseudo-bulk expression across genotypes. Boxed in red are common neuronal senescence markers like p21 (CDKN1A), p16 (CDKN2A), etc.; boxed in blue are the senescence-associated secretory phenotype (SASP) genes.

(I) Dot plot of ferroptosis markers pseudo-bulk expression across genotypes.
